# Pantothenate Kinase 4 Governs Lens Epithelial Fibrosis by Negatively Regulating Pyruvate Kinase M2-Related Glycolysis

**DOI:** 10.1101/2022.08.02.502446

**Authors:** Xue Li, Lin-Lin Luo, Rui-Feng Li, Chun-Lin Chen, Min Sun, Sen Lin

**Affiliations:** Department of Ophthalmology, Daping Hospital, Army Medical Center of PLA, Army Medical University, Chongqing, 400042; Department of Neurology, Xinqiao Hospital, The Second Affiliated Hospital, Army Medical University, Chongqing, 400037

**Keywords:** PANK4, Fibrosis, Lens epithelial cell, Epithelial-Mesenchymal Transition, PKM2, Glycolysis, HIF

## Abstract

Lens fibrosis is one of the leading causes of cataract in the elderly population. The primary energy substrate of the lens is glucose from the aqueous humor, and the transparency of mature lens epithelial cells (LECs) is dependent on glycolysis for ATP. Therefore, the deconstruction of reprogramming of glycolytic metabolism can contribute to further understanding of LEC epithelial–mesenchymal transition (EMT). In the present study, we found a novel pantothenate kinase 4 (PANK4)-related glycolytic mechanism that regulates LEC EMT. The PANK4 level was correlated with aging in cataract patients and mice. Loss of function of PANK4 significantly contributed to alleviating LEC EMT by upregulating pyruvate kinase M2 isozyme (PKM2), which was phosphorylated at Y105, thus switching oxidative phosphorylation to glycolysis. However, PKM2 regulation did not affect PANK4, demonstrating the downstream role of PKM2. Inhibition of PKM2 in *Pank4*^−/−^ mice caused lens fibrosis, which supports the finding that the PANK4–PKM2 axis is required for LEC EMT. Glycolytic metabolism-governed hypoxia inducible factor (HIF) signaling is involved in PANK4–PKM2-related downstream signaling. However, HIF-1α elevation was independent of PKM2 (S37) but PKM2 (Y105) when PANK4 was deleted, which demonstrated that PKM2 and HIF-1α were not involved in a classic positive feedback loop. Collectively, these results indicate a PANK4-related glycolysis switch that may contribute to HIF-1 stabilization and PKM2 phosphorylation at Y105 and inhibit LEC EMT. The mechanism elucidation in our study may also shed light on fibrosis treatments for other organs.

## Introduction

Lens fibrosis, which is characterized by hyperproliferation, migration, and epithelial–mesenchymal transition (EMT) of lens epithelial cells (LECs), causes anterior subcapsular cataract (ASC) and posterior capsule opacification (PCO), damaging vision by changing light scattering[1]. Because of the unique lens biological properties, lens fibrosis can act as an ideal model for investigating the molecular mechanisms of tissue and organ fibrosis, such as liver cirrhosis and idiopathic pulmonary and renal fibrosis, which account for more than 45% mortality[2, 3]. EMT is a major cause of the progression of organ and ocular fibrosis, including the eye diseases PCO, ASC, pterygium, and glaucoma, and is a part of wound healing after ocular surgery[4–9]. In fibrotic cataract, the lens epithelium not only loses its integrity and normal function but also is disorganized with excessive fibronectin and collagen in the extracellular matrix. Lens epithelium EMT is regulated by various mechanisms with the classic prominent roles of transforming growth factor (TGFβ)/Smad signaling[10–13] and glycolytic metabolism[14]. The lens is an avascular tissue relying on limited supply of oxygen and glucose[15], and the lens epithelium can utilize oxidative phosphorylation (OXPHOS) for ATP generation by mitochondria; however, the lens fiber cells are reliant on glycolysis to support their energy demands[14]. Hence, the regulation of glycolytic metabolism of LECs is a potential strategy to prevent lens fibrosis. However, findings on the relationship of cataract to statins (HMG-CoA reductase inhibitors) have been inconsistent[16–18]. Acetyl-CoA is the precursor of HMG-CoA, a CoA product produced during the second step of aerobic cellular respiration and pyruvate decarboxylation, which occurs in the mitochondrial matrix. Pantothenate kinase (PANK) 1, PANK2, and PANK3 are key regulatory enzymes involved in the CoA biosynthetic pathway[19], and catalyze the phosphorylation of the vitamin pantothenate to 4′-phosphopantothenate, subsequently leading to CoA formation. However, PANK4, the fourth member of the PANK family, lacks enzyme activity because an important catalytic glutamate site is missing, and it is thus a peculiarity[19–21]. In a previous study, we found a PANK4 mutation in congenital cataract formation[22], which results in LEC apoptosis and the aberrant expression patterns of crystallin. However, the function and mechanism of PANK4 are still under consideration.

Studies in Han Chinese have shown that gene polymorphisms in PANK4 are associated with type 2 diabetes[23], and functional experiments have shown that PANK4 can affect the apoptosis of pancreatic β cells[24]. In diabetic cataract formation, aldose reductase (AR)-mediated polyol accumulation has been shown to lead to EMT, identified by osmotic imbalance and oxidative stress that result in fiber cell swelling, liquefaction, and eventually cataract development[25, 26] [27]. Eventually, AR leads to pyruvate formation, which is limited by pyruvate kinase M2 isozyme (PKM2) in the last step of glycolysis[28, 29]. Rat Pank4 is associated with PKM2 of KOG2201 and KOG4584 domains under both *in vitro* and *in vivo* conditions[30]. PKM2 is expressed in fetal tissues and cancers, and is a part of EMT in human colorectal cancer cells[31]. Its expression is enhanced in cancers and favors the glycolytic pathway and the metastatic potential such as in pancreatic ductal adenocarcinoma tissues and cell lines[32]. PKM2-mediated glycolysis is also involved in inflammasome activation by modulating EIF2AK2 phosphorylation in macrophages[33]. Moreover, PKM2 activation is a potential treatment against diabetic nephropathy by increasing the glucose metabolic flux and inhibiting the production of toxic glucose metabolites[34]. PKM2 is recruited by phosphorylating histone H3 at T11 that results in H3-K9 acetylation and transcription of glycolytic genes, including *SLC2A1*, *MYC*, *CCND1*, and lactate dehydrogenase A, which enhance the Warburg effect, whereas PKM2 is a Cdc25A substrate that forms a positive feedback loop[35–38]. Unlike many other organs, the eye lens grows throughout life, although growth is slower in adults than in the young, depending on glycolytic activities[39]. However, it is unclear whether PKM2-dependent glycolytic metabolism is one of the major activities in lens epithelium maintenance and EMT. Nonetheless, the relationship between PANK4 and PKM2 in cataract and lens epithelium metabolism remains unclear.

In the present study, we investigated the role of PANK4 in lens epithlium EMT and fibrosis. We show that PANK4 was specifically expressed in human and mice lens epithelium, as well as negatively regulated LEC EMT and functioned in a PKM2 phosphorylation-dependent glycolytic metabolism manner. Without PANK4, age-related or injury-related lens epithelium fibrosis was attenuated. The glycolytic metabolism balance was also perturbed. The suppression of the metabolic switch in LEC by PKM2 activation or by PANK4 loss of function promoted LEC EMT. In contrast, the preemptive amplification of the metabolic response by the inhibition of PKM2 promoted lens epithlium EMT, which sheds light on the potential of clinical drugs in preventing PCO.

## Materials and Methods

### Pank4 transgenic mice generation

*Pank4*^−/−^ mice were generated by Nanjing Biomedical Research Institute of Nanjing University, China, as previously described, with 5-bp mutation, in a C57BL/6J background[40]. Briefly, *pank4* knockout mice were generated using the CRISPR/Cas9 system. Cas9 mRNA and sgRNA were co-injected into zygotes. sgRNA direct Cas9 endonuclease caused a cleavage in exon 3 creating a double-strand break. Such breaks were repaired by non-homologous end joining, and resulted in the destruction of *Pank4* gene. The Pank4 targeting sequence was *gacgtggagcaggatcatgagccaccctatgagatctc agtccaggagagatcacagctcgcctgcatttcatcaagtttgagaatacctacatggaagcctgcctggacttcatcag agaccacctagtcaacactgagaccaaggtcatccaggccactgggggtggagcctacaagttcaaggacctcatcg aggagaagctgcgtctgaa*. Offspring were screened for correct genotype by PCR analysis of tail DNA. Mice were housed in cages in a room with a 12-h light-dark cycle with *ad libitum* access to water and rodent chow diet. Animal protocols were approved by the Institutional Animal Care and Use Committee (IACUC) of Army Medical Center, Army Medical University (No. AMUWEC20201055).

### Lens epithelial tissue collection from cataract patients

A total of 73 cataract patients (aged 17 to 89 yrs) with unilateral cataract presenting to the Department of Ophthalmology, Army Medical Center, Army Medical University, participated in this study. In particular, 17-year-old patients suffered from traumatic cataract, while other patients had age-related cataract (ARC). All patients provided informed consent to participate in the study, and all samples were collected with written informed consent from participants. The study protocol was approved by the Ethics Committee of the Army Medical Center (No. 2018-31-32 and No. 2020-131). The inclusion criteria of ARC included: (1) age more than 50 years; (2) no history of congenital cataract, secondary cataracts, or any other eye diseases; (3) no history of systemic diseases, such as autoimmune disease and cancer, chronic inflammatory diseases, and kidney or hepatic diseases; (4) ARC was considered the presence of any type of ARC (including C-ARC, N-ARC, and PS-ARC) in either eye; (5) ARC recorded using the Lens Opacities Classification System III (LOCS III). All procedures involving human material were performed in accordance with the ethical standards of the institutional and national research committee and the Declaration of Helsinki. All patients underwent detailed examination that included photography and slit-lamp microscopy of the lens. Lens epithelium samples were only from healthy patients. Abnormal coagulation function and etiological examinations for hepatitis B, hepatitis C, syphilis, human immunodeficiency virus, diabetes, hypertension, and hyperlipidemia were excluded from the study. The lens epithelium was dissected and delaminated from the lens of patients by a surgeon. After immediately rinsing with sterile saline solution, some samples were transferred to dry ice for further molecular biological experiments, while the other samples were transferred to 4% paraformaldehyde (PFA) for further immunostaining experiments.

### Cell culture and EMT induction

The human lens epithelial cell line (SRA01/04) was presented by Hao-Tian Lin from the Zhong Shan Ophthalmic Center, Sun Yat-Sen University, as previously described[41]. SRA01/04 cells were grown in Dulbecco’s modified Eagle medium (DMEM) supplemented with 10% (v/v) fetal bovine serum, 100 U/mL streptomycin, and 100 U/mL penicillin, and grown at 37°C in a humidified incubator with 5% CO_2_. LECs were seeded in 12-well cell culture plates at 5.0 × 10^4^ cells/mL and incubated in an incubator (37°C and 5% CO_2_) in the standard medium for 24 h. LECs were treated with TGF-β2 (1 μg/mL) for 12 h until the cell density increased to 60%. Next, LECs were divided into groups to deal with PKM-IN-1 or DASA-58 separately for 24 h.

### Lens culture and treatment

All experimental procedures conformed to the Association for Research in Vision and Ophthalmology (ARVO) Statement for the Use of Animals in Ophthalmic and Vision Research. Briefly, the lens culture protocol followed a previous report with some modification[42–44]. Briefly, lenses from one-month C57BL/6J mice were carefully removed using a forceps and a pair of fine scissors, and maintained in 4 mL serum-free DMEM/F12 medium, containing 0.1 mg/mL L-glutamine, 50 IU/mL penicillin, and 50 mg/mL streptomycin. TGF-β2 was added to the culture medium at a final concentration of 5 ng/mL. The culture medium was renewed every second day throughout the culture period. Lenses were cultured for up to 5 days, and the images were captured using a dissecting microscope (Olympus, Tokyo, Japan).

### Immunofluorescence staining

For mice, the eyeballs were removed and fixed in 4% PFA for 3 h, and dehydrated in a sucrose solution (30%) for 3 h. Eyeballs were embedded in OCT Tissue Tek Medium, and cut into 10-µm-thick sections adhered onto anti-slip slides using a frozen section machine. LECs were grown on 13-mm coverslips in 24-well plates and fixed in 4% PFA for 15 min at room temperature (RT). Eyeball sections and fixed LECs were washed with phosphate-buffered saline (PBS), and samples were permeabilized and blocked in 0.3% Triton X-100 and 3% bovine serum albumin in PBS for 30 min at RT. Samples were incubated with appropriate primary antibodies (PANK4 was purchased from Cell Signaling Technology (CST), Cat.12055, rabbit anti-mouse, 1:100 for IF; PKM2 from CST, Cat. 4053, rabbit anti-mouse, 1:100 for IF; PKM2 from Bioss, Cat. bs-0102M, mouse anti-mouse, 1:200 for IF; p-PKM2^Y105^ from Abcam, Cat. ab156856, 1:100 for IF; p-PKM2^ser37^ from Thermo Fisher Scientific, Cat. PA5-37684, 1:200 for IF) (Table 1) overnight at 4°C. PANK4 antibody specificity was identified in tissues from genetic knockout, heterozygous and WT mice from the same littermate as described previously [40]. After these steps, the samples were washed with PBS and incubated with fluorescent secondary antibodies (Alexa Fluor® 594-AffiniPure Goat Anti-Rabbit IgG (H+L), Cat. 111-585-003, Alexa Fluor® 488-AffiniPure Goat Anti-Rabbit IgG (H+L), Cat. 111-545-003 and Alexa Fluor® 594-AffiniPure sheep Anti-mouse IgG (H+L), Cat. 515-585-003 were purchased from Jackson ImmunoResearch, 1:500 for 2^nd^ antibodies conjugations) (Table 1) for 0.5 h at RT. Samples were mounted after DAPI staining. In this process, samples were washed with PBS three times for 3 min between each step. Isotype antibody controls (Rabbit IgG, polyclonal - Isotype Control, Cat. ab172730 and mouse IgG2b, kappa monoclonal [7E10G10] - Isotype Control, Cat. ab170192 were purchased from Abcam) and secondary antibody only controls were also performed to validate antibody specificity. Images were captured with a confocal microscope (Olympus FV3000, Japan).

### Protein extraction and Western blot

The protein was obtained from lens tissues in *in vivo* experiments or primary cultured SRA01/04 cells *in vitro*. Samples were lysed in ice-cold lysis buffer, which consisted of RIPA buffer and the protease inhibitor cocktail. The protein was then clarified by Heraeus Sepatech, and the supernatant was collected in new tubes. The protein concentration was determined using the bicinchoninic acid (BCA) protein assay kit. The loading buffer was added to protein, and the mixture was boiled for 10 min and stored at −80°C.

For Western blot, 20 µg of total protein was used to separate on 10% sodium dodecyl sulfate (SDS)-polyacrylamide gels, and transferred to polyvinylidene fluoride membranes. The membranes were blocked for 2 h in Tris-buffered saline with 5% non-fat skimmed milk powder at RT. The membranes were then probed with primary antibodies (PKM1, Cat. 7067; PKM2, Cat. 4053; N-cadherin, Cat. 14215S; α-SMA, Cat. 48938; HIF-1α, Cat. 36169; HIF-2α, Cat. 59973; HIF-1β, Cat. 5537 and GAPDH Cat. 5174 were all rabbit anti-mouse antibodies and purchased from CST, with working concentration of 1:1000. p-PKM2^Y105^ from Abcam, Cat. ab156856, 1:1000 for WB; p-PKM2^ser37^ from Thermofisher, Cat. PA5-37684, 1:2000 for WB) (Table 1) at 4°C overnight and probed by the secondary antibodies at RT for 1.5 h. Finally, the images were captured by the Western blot automatic exposure apparatus.

### Co-immunoprecipitation (Co-IP)

Co-immunoprecipitation was performed as described in the manual. Briefly, after thorough resuspension, magnetization, and washing, SureBeads (Biorad, 1614013) were resuspended with 5 µg PKM1 (CST, D30G6, Cat. No. 7067), PKM2 (CST, D78A4, Cat. No. 4053), and PANK4 (CST, D8N2C, Cat. No. 12055) antibodies in a final volume of 200 µl, respectively, and IgG as the control. Rotate for 10 min at RT. Supernatant was discarded, and the beads were washed three times with 1,000 µl PBS-T. Total lysate (300 μg) from WT and *Pank4*^−/−^ mice lens epithelium was diluted to a final volume of 500 μL NP40 lysis buffer (Solarbio, N8032), and phenylmethylsulfonyl fluoride (PMSF, Solarbio, P0100). Rotate for 1 h at RT. Magnetized beads were removed and the supernatant was discarded. The beads were washed with 1,000 µl PBS-T, resuspend the beads thoroughly, centrifugated, magnetized beads and supernatant was discarded; this process was repeated three times. Before the last magnetization, the resuspended beads were transferred to a new tube. All tubes were spun down for several seconds. Magnetized beads were removed, and the residual buffer was aspirated from the tubes, and 40 µl 1× Laemmli buffer (biorad,1610747) was added and incubated for 10 min at 70°C. Magnetized beads were removed, and the eluent was moved to a new vial. Equal amounts of extracts were separated by SDS-PAGE, and then they were transferred onto nitrocellulose membranes and then blotted with specific antibodies.

### Wound healing assay

SRA01/04 cells were cultured in a 12-well plate. After cells were grouped and received the corresponding treatment. When cell density increased to 90%, a 10-μL pipette tip was used to make a straight scratch through the monolayers. In the following period, cells migrated to the wound surface. The scratch width was used to represent the migration ability of SRA01/04 cells. After 24 h, the scratch width was measured in five randomly selected fields. Data presented are means of triplicates.

### ATP measurement

Intracellular ATP was determined using an ATP assay kit as previously described[45]. Briefly, cells were lysed for 5 min and centrifuged at 12,000 g at 4°C for 5 min. The supernatant was added to test plates pre-treated with the ATP detection reagent. Luminescence was measured using a microplate reader. The remaining cell lysate was measured with the BCA protein assay kit to determine the protein concentration. Finally, the ATP concentration was expressed as μmol/g protein.

### mRNA sequencing and bioinformatics data analysis

#### Total RNA extraction, mRNA library construction, and RNA sequencing (RNA-seq)

Total RNA was extracted from the lens epithelial tissues using TRIzol (Invitrogen, Carlsbad, CA, USA) according to previously described protocols[46, 47]. Briefly, after routine RNA extraction, 25–100 µL of diethyl pyrocarbonate-treated water was added to dissolve RNA. First-strand cDNA was generated using random hexamer-primed reverse transcription, followed by second-strand cDNA synthesis. A-tailing mix and RNA index adapters were added for incubating for end repair. cDNA fragments were amplified using PCR, and the products were purified using Ampure XP beads (Beckman Coulter) dissolved in EB solution. The product was confirmed using an Agilent 2100 Bioanalyzer for quality control. The double-stranded PCR products were heated, denatured, and circularized by the splint oligo sequence to derive the final library. The single-stranded circular DNA was formatted as the final library. The final library was amplified with phi29 to prepare a DNA nanoball (DNB), with more than 300 copies. DNBs were loaded into the patterned nanoarray, and single-end 50-base reads were generated on an MGISEQ-2000 sequencing platform. mRNA sequencing was performed by Genetics Biotech Inc. and BioWavelet Ltd. in Chongqing, China.

### Identification of differentially expressed genes

The expression level of genes was calculated using Salmon (v1.4.0). The heatmap was drawn using pheatmap (v1.0.8). Essentially, differential expression analysis was performed using DESeq2 (v1.4.5) with Q-value ≤0.05. To gain insight into the change in phenotype, Gene Ontology (http://www.geneontology.org/) and Kyoto Encyclopedia of Genes and Genomes (https://www.kegg.jp/) enrichment analyses of the annotated differentially expressed genes were performed using pHYPER based on the hypergeometric distribution. The significant terms and pathways were corrected with Q-value with a rigorous threshold (Q-value ≤0.05) using the Bonferroni method.

### Pyruvate detection

The PA levels were determined by the pyruvate assay kit (Pyruvate Assay Kit (ab65342/K609-100), Abcam). The PA content was evaluated by measuring the absorbance at 505 nm, and the PA content was calculated using the following formula (A is absorbance): PA content (µmol/mL) = (A (sample) − A (blank)) / (A (standard) − A (blank)) × standard concentration (0.2 µmol/mL) / protein concentration of sample (mg/mL).

### Seahorse measurement

Seahorse analysis was performed as previously described[48]. The oxygen consumption rate (OCR) and extracellular acidification rate (ECAR) were determined using the Seahorse XF96 Extracellular Flux Analyzer through the Seahorse XF Mito Stress Test Kit and Seahorse XF Glycolysis Stress Test Kit. It is used to measure cell dependence on oxidative metabolism or preferentially the glycolytic function in cells. LECs were counted and seeded in XF96 Seahorse® microplates with 10,000 cells per well. After cells were grouped and received the corresponding treatment. When cell density reached 80%, cells were cultured in XF assay medium supplemented with 1 mM glutamine to assess ECAR and in XF assay medium supplemented with 2 mM glutamine to assess OCR. The cell cultures were placed in a CO_2_-free incubator with 37°C for 0.5 h. The plate was left to equilibrate in before being transferred to the Seahorse XF96 analyzer. The pre-hydrated cartridge was filled with the indicated compounds and calibrated for 30 min in the Seahorse Analyzer. All experiments were performed at 37°C. Normalization to protein content was performed after each experiment. The Seahorse XF Report Generator automatically calculated the parameters from wave data that were exported to GraphPad Prism software (v8.4, GraphPad Software).

### Statistical analysis

Statistical analysis was performed using GraphPad Prism 8.0 software. Data are presented as means ± SD. Shapiro-Wilk normality test was performed to assess the normality of the distribution of data. Due to the small sample sizes, we acknowledge the assessment of normality may not reliable. For comparison between two groups, Student’s *t*-test was performed. For comparisons among more than two groups, one-way analysis of variance followed by Bonferroni’s post hoc test was performed. All included data were tested for normality. If normal distribution was not achieved, a nonparametric test was used. Non-parametric comparisons (Mann-Whitney U Test or Kruskal-Wallis Test) were used for non-normally distributed data or n < 6. A *P* value < 0.05 was considered statistically significant.

## Results

### PANK4 levels increase with age in both rodent and human lens epithelial tissues

PCO-related fibrosis is reported to be associated with lens epithelium aging[12]. In this study, PANK4-positive punctate structures were specifically located in the cytoplasm of LECs of 17-, 47-, and 81-year-old patients (Fig. 1A). The fluorescence intensity of 47- and 81-year-old groups exhibited more intensive gray value in the cytoplasm (Fig. 1B). Furthermore, an increased number of puncta was found in the intracellular cytoplasm of LECs in the aged groups (47 yrs and 81 yrs, respectively) compared with that in younger groups (Fig. 1C; *P* < 0.0001, n = 19). To verify the immune-intensity results, we collected and pooled lens epithelium from patients aged 50–60 yrs, 60–70 yrs, 70–80 yrs, and over 80 yrs groups. The intensities of PANK4 blots were significantly elevated in 70–80 yrs and over 80 yrs female groups compared with the 50–60 yrs female group, respectively (Fig. 1D; *P*=0.384 *vs* 60-70yrs group, n=6; *P*=0.0073 *vs* 70-80yrs group, n=6; *P*=0.0028 *vs* 80yrs+ group, n=6). Similarly, the intensities of PANK4 blots were significantly elevated in 70–80 yrs and over 80 yrs male groups compared with the 50–60 yrs group, respectively (Fig. 1E; *P*=0.1514 *vs.* 60–70 yrs group, n=6; *P*=0.0096 *vs.* 70–80 yrs group, n=6; *P* < 0.0001 *vs.* 80 yrs + group, n=6). In mice, PANK4 was localized in the whole anterior lens epithelium and equator by immunostaining (Fig. 1F). Amplified high power crops showed PANK4-positive signals in cytoplasmic localization (Fig. 1F inner panel), which were consistent with immunostaining, suggesting the promising pattern of PANK4 in the lens. PANK4 also increased dramatically in 2 Mo, 6 Mo, and 12 Mo groups rather than in the P15 group (Fig. 1G; *P*=0.0007 *vs.* 2 Mo group, n=6; *P*=0.011 *vs.* 6 Mo group, n=6; *P*=0.0003 *vs.* 80 yrs+ group, n=6), which strengthen the results of PANK4 in humans. Meanwhile, the lens epithelium EMT-related characteristic molecule, such as N-cadherin, was significantly upregulated in 6 Mo and 12 Mo groups compared with the P15 group (Fig. 1I, N-cad panel, *P*=0.0317 *vs.* 2 Mo group, n=6; *P*<0.0001 *vs.* 6 Mo group, n=6; *P*<0.0001 *vs.* 12 Mo group, n=6). The collagen I level was increased in the 2 Mo, 6 Mo, and 12 Mo groups compared with the P15 group (Fig. 1I, Col I panel, *P*=0.0361 *vs.* 2 Mo group, n=6; *P*=0.0018 *vs.* 6 Mo group, n=6; *P*=0.0005 *vs.* 12 Mo group, n=6). The α-SMA level was elevated in 2 Mo and 12 Mo groups compared with the P15 group. No significant difference was found in the 6 Mo group compared with the P15 group (Fig. 1I; α-SMA panel, *P*=0.0218 *vs.* 2 Mo group, n=6; *P*=0.0088 *vs.* 6 Mo group, n=6; *P*=0.0005 *vs.* 12 Mo group, n=6). Thus, PANK4 was correlated with age and age-dependent lens epithelium fibrosis.

**Figure 1.**
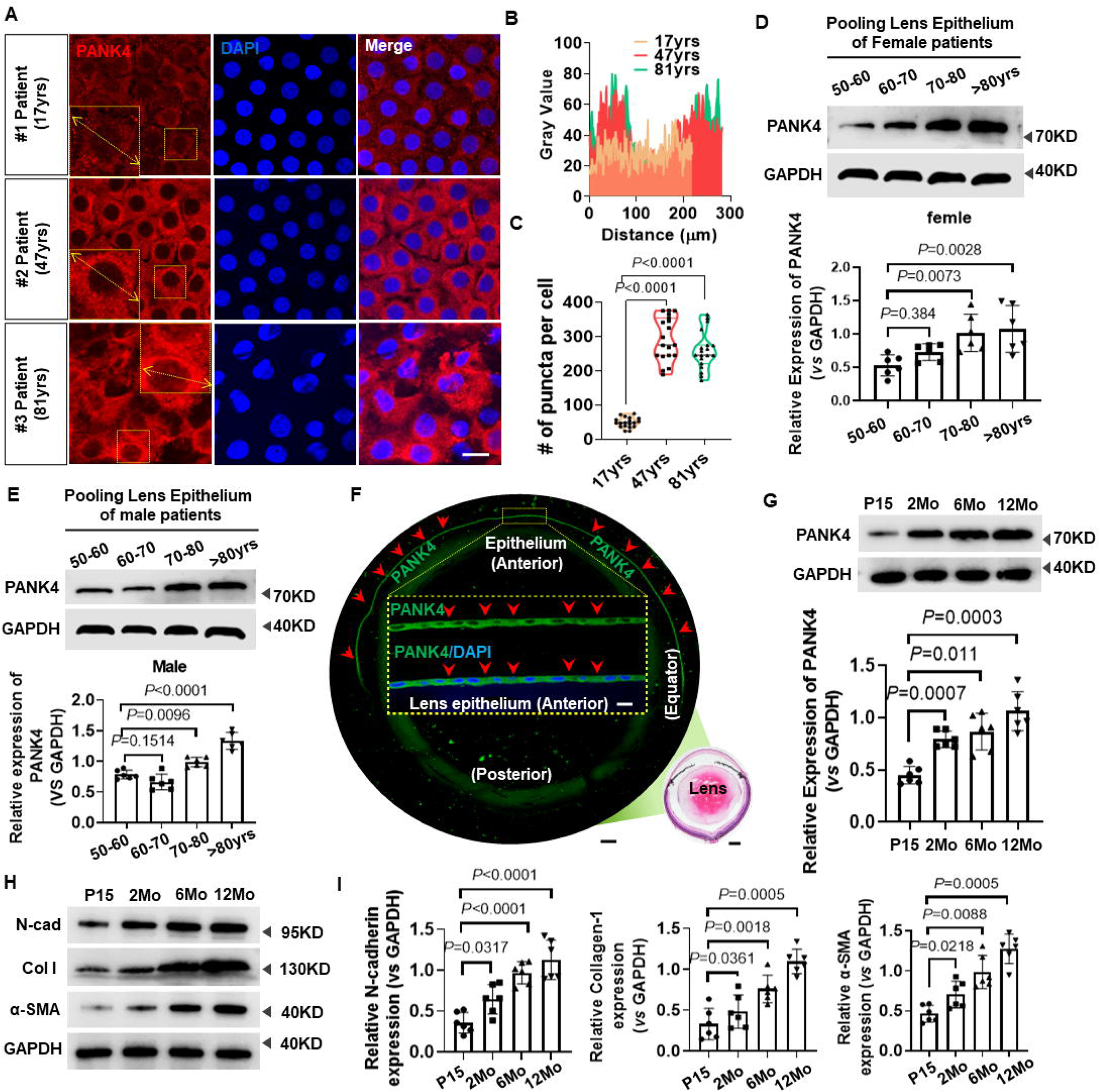
PANK4 elevated with age and related to lens fibrosis and EMT. (**A**) Immunostaining of PANK4 in lens epithelium from patients aged at 17yrs, 47yrs and 81yrs. The scale bar = 10μm. (**B**) Plot profile analysis of immunostaining in lens epithelium of patients aged from 17yrs, 47yrs and 81yrs. (**C**) Quantification of PANK4 immunostaining positive puncta per cell in LECs, from patients aged at 17yrs, 47yrs and 81yrs. n=19, *P*<0.0001. (**D**) The levels of PANK4 protein in lens epithelium from female cataract patients (50-60yrs, 60-70yrs, 70-80yrs and more than 80yrs groups) were determined by Western blotting assay and quantified by densitometry. Data were analyzed by one way ANOVA plus Bonferroni post hoc test. All data shown are mean ± SD. Relative expression is the fold changes relative to the 50-60yrs group. *P*=0.384 *vs* 60-70yrs group, n=6; *P*=0.0073 *vs* 70-80yrs group, n=6; *P*=0.0028 *vs* 80yrs+ group, n=6. (**E**) The levels of PANK4 protein in lens epithelium from male cataract patients (50-60yrs, 60-70yrs, 70-80yrs and more than 80yrs groups) were determined by Western blotting assay and quantified by densitometry. Data were analyzed by one way ANOVA plus Bonferroni post hoc test. All data shown are mean ± SD. Relative expression is the fold changes relative to the 50-60yrs group. *P*=0.1514 *vs* 60-70yrs group, n=6; *P*=0.0096 *vs* 70-80yrs group, n=6; *P*<0.0001 *vs* 80yrs+ group, n=6. (**F**) Immunostaining pattern of PANK4 in the lens epithelium of adult mice. The scale bar = 200μm. HE image presented an anatomy structure of lens. Inner panel: High power images of PANK4 immunostaining results in anterior lens epithelium and equator of adult WT mice. The scale bar = 10μm. (**G**) The levels of PANK4 protein in lens epithelium of from P15, 2Mo, 6Mo and 12Mo mice, detected by Western blotting assay and quantified by densitometry. Data were analyzed by one way ANOVA plus Bonferroni post hoc test. All data shown are mean ± SD. Relative expression is the fold changes relative to the P15 group. *P*=0.0007 vs 2Mo group, n=6; *P*=0.011 vs 6Mo group, n=6; *P*=0.0003 vs 80yrs+ group, n=6. (**H** and **I**) The levels of N-cadherin, collagen I and α-SMA protein in lens epithelium of from P15, 2Mo, 6Mo and 12Mo mice, detected by Western blotting assay and quantified by densitometry. Data were analyzed by one way ANOVA plus Bonferroni post hoc test. Relative expression is the fold changes relative to the P15 group. Results are expressed as means ± SD. N-cad panel, *P*=0.0317 *vs* 2Mo group, n=6; *P*<0.0001 *vs* 6Mo group, n=6; *P*<0.0001 *vs* 12Mo group, n=6. Col I panel, *P*=0.0361 *vs* 2Mo group, n=6; *P*=0.0018 *vs* 6Mo group, n=6; *P*=0.0005 *vs* 12Mo group, n=6. α-SMA panel, *P*=0.0218 *vs* 2Mo group, n=6; *P*=0.0088 *vs* 6Mo group, n=6; *P*=0.0005 *vs* 12Mo group, n=6.

### PANK4 is required for lens epithelium fibrosis and EMT

We next considered whether PANK4 modification alters EMT. Dramatic decrease was observed in levels of N-cadherin (*P*=0.0065, n=6), collagen I (*P*=0.0026, n=6), and α-SMA (*P*=0.005, n=6) in the lens epithelium of 2-Mo *Pank4*^−/−^ mice (Fig. 2A and B). To determine whether PANK4 is required for age-related lens fibrosis, PANK4 expression was analyzed in 1-year-old WT and *Pank4*^−/−^ mice. Similarly, N-cadherin (*P*=0.0198, n=6), collagen I (*P*=0.038, n=6), and α-SMA (*P*=0.0008, n=6) were also inhibited when PANK4 was deficient (Fig. 2C and D). TGF-β2, the major isoform in the aqueous humor, acts as a predominant pathogenic factor in the development of lens epithlium EMT[49, 50]. To determine the effect of loss of function of PANK4 in EMT alleviation, we next established a lens fibrosis model by TGF-β2 injection into mice eyes. Compared with the WT group, *Pank4*^−/−^ mice showed a dramatic decrease in N-cadherin (*P*=0.0422 in WT vs WT+TGFβ2 group, n=6; *P*=0.0075 in WT vs *Pank4*^−/−^ group, n=6; *P*<0.0001 in *Pank4*^−/−^ vs *Pank4*^−/−^+ TGFβ2 group, n=6), collagen I (*P*=0.0189 in WT vs WT+TGFβ2 group, n=6; *P*=0.0336, in WT vs *Pank4*^−/−^ group, n=6; *P*<0.0001 in *Pank4*^−/−^ vs *Pank4*^−/−^+ TGFβ2 group, n=6), and α-SMA (*P*<0.0011 in WT vs WT+TGFβ2 group, n=6; *P*=0.0242, in WT vs *Pank4*^−/−^ group, n=6; *P<*0.0001 in *Pank4*^−/−^ vs *Pank4*^−/−^+ TGFβ2 group, n=6) levels with and without TGF-β2 injection (Fig. 2E and F). The alleviation of lens fibrosis was either found in TGF-β2-treated groups or lens puncture groups (Fig. 2G) compared with WT mice. Thus, PANK4 was required for lens epithlium EMT. To confirm the effect of loss of function of PANK4, siRNA*^Pank4^* was used to knock it down in SRA01/04 cells. Here, we assessed three siRNA sequences—#274, #962, and #1114, of which #1114 showed almost 80% inhibition of PANK4, whereas the other two siRNAs exhibited attenuated inhibitory effect of nearly 30% (Fig. S1A). After 48-h interference of *Pank4* mRNA, the collagen I (*P*=0.0038 in NC vs siPank4 group, n=6; *P*=0.0048 in siControl vs siPank4 group, n=6), N-cadherin (*P*=0.0381 in NC vs siPank4 group, n=6; *P*=0.0369 in siControl vs siPank4 group, n=6) and α-SMA (*P*<0.0001 in NC vs siPank4 group, n=6; *P*=0.0002 in siControl vs siPank4 group, n=6) levels remarkably decreased (Fig. 2H and I). These results were consistent with the decrease in the SRA01/04 cell migration test (Fig. 2J and K) (*P*<0.0001 in NC vs siPank4 group, n=7; *P*<0.0001 in siControl vs siPank4 group, n=7; 24h group), in which siRNA*^Pank4^* inhibited the migration rate of SRA01/04 cells. PANK4 overexpression by Lentivirus–*Pank4 (Pank4* OE) accelerated the migration (Fig. 2J and K) (*P*<0.0001 in NC vs *Pank4* OE group, n=7; *P*<0.0001 in siControl vs *Pank4* OE group, n=7; 12h and 24h groups, respectively), which suggested that PANK4 was implicated in LEC EMT.

**Figure 2.**
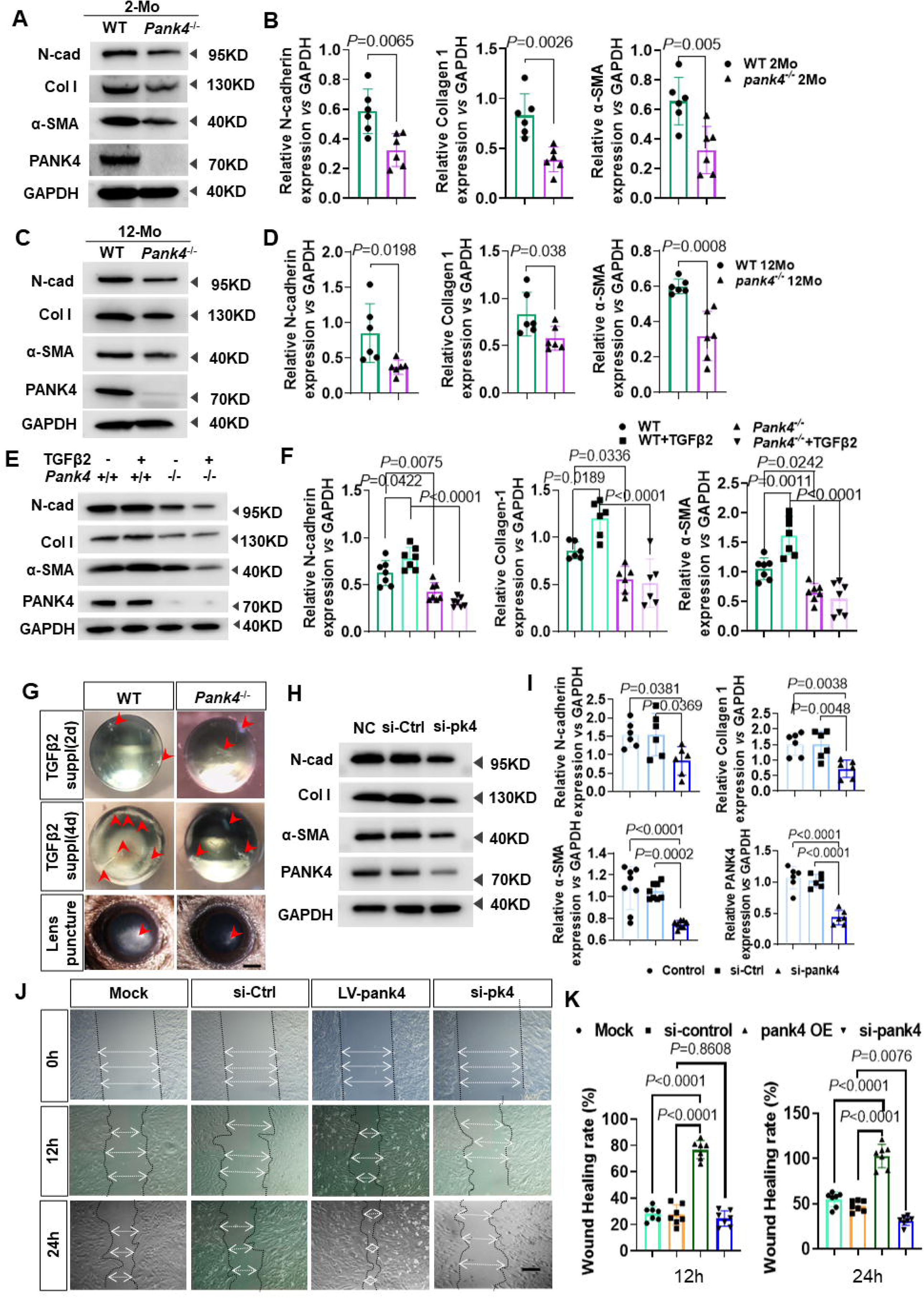
PANK4 is required for lens epithelium fibrosis and EMT. (**A**) and (**B**) The levels and quantifications of N-cadherin, collagen I and α-SMA protein in 2-month-old (2Mo) WT and *Pank4*^−/−^ lens epithelium. anti-PANK4 as positive control. Data were analyzed by student’s *t* test. Relative expression is the fold changes relative to the 2Mo WT group. N-cadherin, *P*=0.0065, n=6; collagen I, *P*=0.0026, n=6; α-SMA, *P*=0.005, n=6. (**C**) and (**D**) The levels and quantifications of N-cadherin, collagen I and α-SMA protein in 12-month-old (12Mo) WT and *Pank4*^−/−^ lens epithelium. anti-PANK4 as positive control. Data were analyzed by student’s *t* test. Relative expression is the fold changes relative to the 12Mo WT group. N-cadherin, *P*=0.0198, n=6; collagen I, *P*=0.038, n=6; α-SMA, *P*=0.0008, n=6. (**E**) and (**F**) The levels and quantifications of N-cadherin, collagen I and α-SMA protein in 2-month-old (2Mo) WT and *Pank4*^−/−^ lens epithelium, with and without TGF-β2 injection. Data were analyzed by one way ANOVA plus Bonferroni post hoc test. Relative expression is the fold changes to comparable groups. N-cadherin: *P*=0.0422 in WT vs WT+TGFβ2 group, n=6; *P*=0.0075 in WT vs *Pank4*^−/−^ group, n=6; *P*<0.0001 in *Pank4*^−/−^ vs *Pank4*^−/−^+ TGFβ2 group, n=4. collagen I: *P*=0.0189 in WT vs WT+TGFβ2 group, n=6; *P*=0.0336, in WT vs *Pank4*^−/−^ group, n=6; *P*<0.0001 in *Pank4*^−/−^ vs *Pank4*^−/−^+ TGFβ2 group, n=6. α-SMA: *P*=0.0011 in WT vs WT+TGFβ2 group, n=6; *P*=0.0242, in WT vs *Pank4*^−/−^ group, n=6; *P<*0.0001 in *Pank4*^−/−^ vs *Pank4*^−/−^+ TGFβ2 group, n=6. (**G**) Cultured lens with TGF-β2 (5.0ng/mL) addition for 2 and 4 days (solid arrow indicated fibrosis in the lens). In vivo lens fibrosis phenotype, 7 days after puncture experiment in WT and *Pank4*^−/−^ mice. Observed and recorded by slit light microscope. (**H**) and (**I**) The levels and quantifications of N-cadherin, collagen I, α-SMA and PANK4 protein after siRNA administration. Data were analyzed by one way ANOVA plus Bonferroni post hoc test. Relative expression is the fold changes relative to the NC group. N-cadherin: *P*=0.0381 in NC vs siPank4 group, n=6; *P*=0.0369 in siControl vs siPank4 group, n=6; Collagen I: *P*=0.0038 in NC vs siPank4 group, n=6; *P*=0.0048 in siControl vs siPank4 group, n=6; α-SMA: *P*<0.0001 in NC vs siPank4 group, n=6; *P*=0.0002 in siControl vs siPank4 group, n=6. PANK4: *P*<0.0001 in NC vs siPank4 group, n=6; *P*<0.0001 in siControl vs siPank4 group, n=6. (**J**) and Lens epithelial cell migration assay at 0h, 12h and 24h, with Lenti virus-*Pank4* overexpression and siRNA interference treatments. (**K**) Quantifications of the wound healing rate (%) at 0h, 12h and 24h. Results are expressed as means ± SD. *P*<0.0001 in NC vs *Pank4* OE group, n=7; *P*<0.0001 in siControl vs *Pank4* OE group, n=7; 12h and 24h groups, respectively.

### PANK4 negatively regulates PKM2 in LECs

PKM2 is considered as an associated protein that interacts with rat PANK4, which is isolated from the rat muscle cDNA library by yeast two-hybrid screening[30]. In addition, by STRING protein–protein interaction networks functional enrichment analysis (https://string-db.org/), we found a potential relationship between PANK4 and PKM with a score of 0.705. The Co-IP blot results showed that PKM2 significantly increased on immunoprecipitation with PANK4, whereas PKM1 did not. To verify their binding relationship, PKM2 antibody was used to bait the protein, which showed significant PANK4 immunoprecipitation (Fig. 3A), indicating the binding relationship between PANK4 and PKM2. It clearly indicated the specific location of PKM2 in the lens epithelium, and the co-staining pattern of PKM2 and PANK4 in the LEC cytoplasm (Fig. 3B). We then investigated their regulatory mechanism using *Pank4*^−/−^ mice, which demonstrated a significantly elevated level of PKM2 in the lens epithelium by immunostaining (Fig. 3C and D) as well as Western blotting results (Fig. 3E and F; PANK4: *P*<0.0001, n=6; PKM2: *P*<0.0001, WT vs *Pank4*^−/−^, n=6), but with no significant difference in PKM1 expression between WT and *Pank4*^−/−^ groups (*P*=0.805, WT *vs. Pank4*^−/−^, n=6). By siRNA*^Pank4^*knockdown of *Pank4* mRNA in SRA01/04, a remarkable increase was observed in PKM2 expression in the cytoplasm (Fig. 3G), which was consistent with the results of the genome deficiency of *Pank4* in the lens epithelium, followed by a Western blot confirmation of the elevated PKM2 protein level (Fig. 3H and I, *P*=0.0002, siCont *vs.* siPank4; *P*<0.0001, NC *vs.* siPank4, n=6) without a significant change in PKM1 (Fig. 3H and I; *P*=0.9742, siCont *vs.* siPank4, n=6). TGF-β2 injection induced lens epithelium fibrosis, and *Pank4* deletion increased the PKM2 level as well (Fig. 3J and K; *P*=0.0069, WT *vs. Pank4*^−/−^; *P*=0.0036, WT+TGFβ *vs. Pank4*^−/−^+TGFβ, n=6), suggesting the negative regulation between PANK4 and PKM2. In the conversion of phosphoenolpyruvate (PEP) to pyruvic acid, PEP is alternatively catalyzed by PKM2[53]. Strikingly, pyruvic acid increased when PANK4 was inhibited in LECs (Fig. 3L; *P* < 0.0001, siControl *vs.* siPank4, n=6). These data suggest a role for the PANK4–PKM2 axis in glycolysis in the lens epithelium (Fig. 3M).

**Figure 3.**
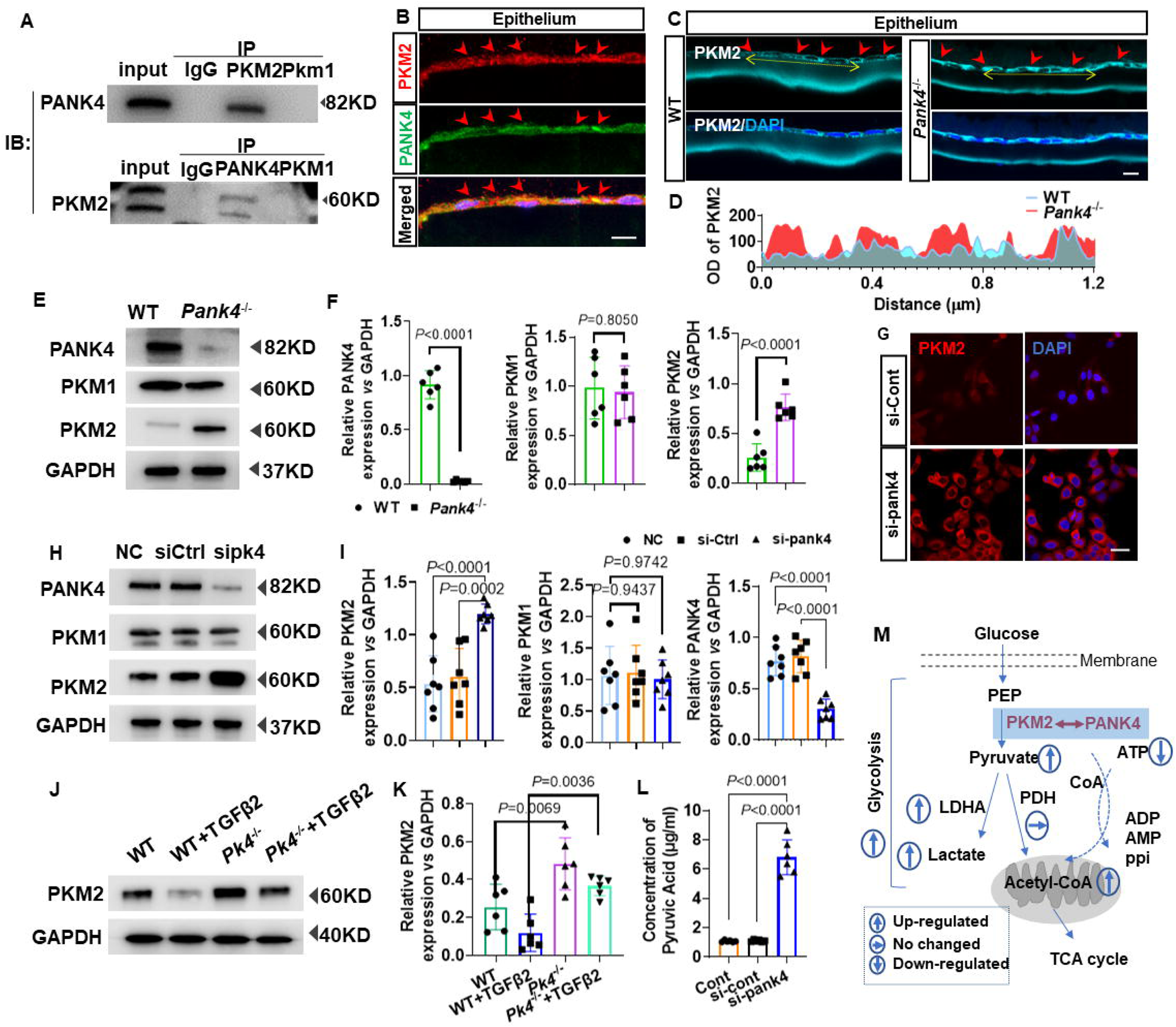
PANK4 negatively regulating PKM2 in lens epithelium. (**A**) Co-IP results of PANK4 and PKM2. (**B**) Co-Immunofluorescence staining of PKM2 and PANK4 in lens epithelium. The scale bar = 10μm. (**C**) Immunofluorescence staining of PKM2 in lens epithelium of WT and *Pank4*^−/−^ lens. The scale bar = 10μm. (**D**) Plot profile curve of PKM2 in WT and *Pank4*^−/−^ lens. (**E**) and (**F**) Protein levels and quantifications of PANK4, PKM1, PKM2 in WT and *Pank4*^−/−^ lens. GAPDH as inner control. Data were analyzed by student’s *t* test. Relative expression is the fold changes relative to the WT group. PANK4: *P*<0.0001, n=6; PKM1: *P*=0.805 PKM2: *P*<0.0001, WT vs *Pank4*^−/−^, n=6. (**G**) Immunofluorescence staining results of PKM2 in LECs in siControl group and si*Pank4* group. The scale bar = 10μm. (**H**) and (**I**) Protein levels and quantifications of PANK4, PKM1, PKM2 in si-control group and siPank4 group. Data were analyzed by one way ANOVA plus Bonferroni post hoc test. Relative expression is the fold changes relative to the siPank4 group. PKM2: *P*<0.0001, NC vs siPank4; *P*=0.0002, siCtrl vs siPank4, n=6. PKM1: *P*=0.9437, NC vs siPank4, n=6; *P*=0.9742, siCtrl vs siPank4, n=6. PANK4: *P*<0.0001, NC vs siPank4; *P*<0.0001, siCtrl vs siPank4, n=6. (**J**) and (**K**) Protein levels and relative expression quantifications of PKM2 in lens epithelium of WT, WT with TGF-β2 injection, *Pank4*^−/−^, *Pank4*^−/−^ with TGF-β2 injection groups. Data were analyzed by one way ANOVA plus Bonferroni post hoc test. Relative expression is the fold changes relative to the comparable groups.*P*=0.0069, WT vs *Pank4*^−/−^; *P*=0.0036, WT+TGFβ vs *Pank4*^−/−^+TGFβ, n=6. (**L**) Concentrations of pyruvic acid (μg/ml) in LECs of negative control group, si-control group and siPank4 group. Data were analyzed by one way ANOVA plus Bonferroni post hoc test. Relative expression is the fold changes relative to the siPank4 group. *P*<0.0001, siControl vs siPank4, n=6; *P*<0.0001, NC vs siPank4, n=6. (**M**) Schematic illustration of current results.

### Inhibition of PKM2 attenuates lens epithelium EMT by promoting PKM2 phosphorylation at Y105 residue

The inhibition and activation of PKM2 by PKM-IN-1 and DASA-58, which are respectively typical antagonist and agonist of PKM2, did not significantly change the PANK4 level, suggesting a downstream role for PKM2. In the presence of PKM2 inhibition or activation by PKM-IN-1 or DASA-58, the N-cadherin levels (*P*=0.0033, Veh *vs.* DASA-58 groups, n=6), collagen I (*P*=0.0002, Veh *vs.* DASA-58 groups, n = 6), and α-SMA (*P*=0.0007, Veh *vs.* DASA-58 groups, n=6) significantly decreased when activated by DASA-58. However, only N-cadherin and α-SMA levels increased when treated by PKM-IN-1. (N-cad: *P*=0.0138, Veh *vs.* PKM-IN-1 groups, n=6; Col-1: *P*=0.1066, Veh *vs.* PKM-IN-1 groups, n=6; α-SMA: *P*=0.0019, Veh *vs.* PKM-IN-1 groups, n=6;) (Fig. 4A and B). These results indicate an important role for PKM2 downstream of PANK4 in regulating LEC EMT.

**Figure 4.**
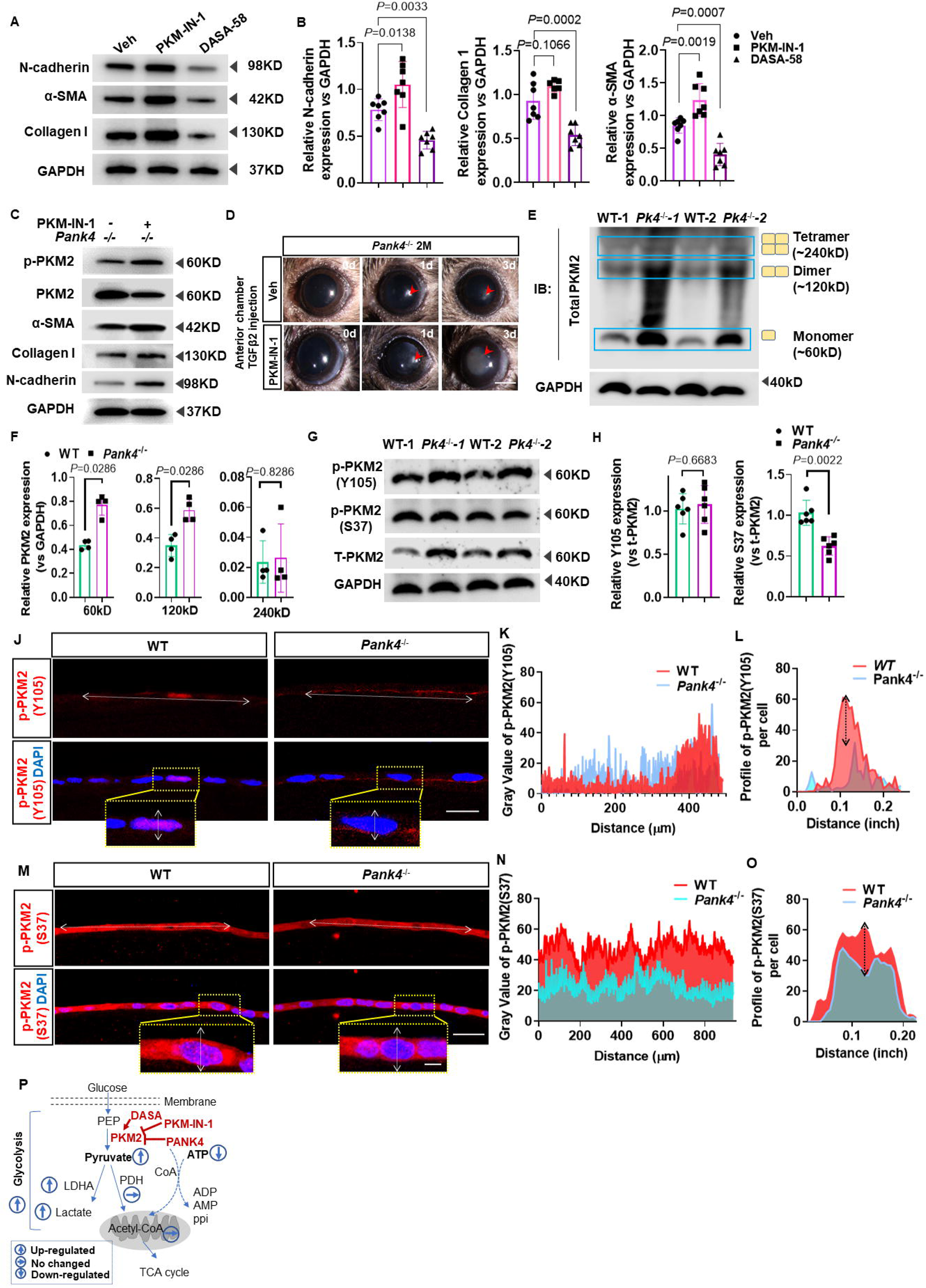
PANK4 deficiency is responsible for PKM2 Y105 phosphorylation. (**A**) and (**B**) Protein levels and quantifications of N-cadherin, α-SMA and collagen-I under PKM-IN-1 and DASA-58 treatments. Data were analyzed by one way ANOVA plus Bonferroni post hoc test. Relative expression is the fold changes relative to the Veh group. N-cadherin: *P*=0.0138, Veh vs PKM-IN-1 groups, n=6; *P*=0.0033, Veh vs DASA-58 groups, n=6; collagen I: *P*=0.1066, Veh vs PKM-IN-1 groups, n=6; *P*=0.0002, Veh vs DASA-58 groups, n=6; α-SMA(*P*=0.0019, Veh vs PKM-IN-1 groups, n=6; *P*=0.0007, Veh vs DASA-58 groups, n=6). GAPDH as inner control. (**C**) Protein levels and quantifications of p-PKM2, PKM2, N-cadherin, α-SMA and collagen-I under PKM-IN-1 inhibition in *Pank4*^−/−^ mice. (**D**) TGF-β2 injection with vehicle and PKM-IN-1 treatment in 2Month old *Pank4*^−/−^ mice. (**E**) Represented blot image of mice lens epithelium samples to show PKM2 monomer, dimer and tetramer. (**F**) Relative expression of t-PKM2 vs GAPDH. Data were analyzed by Mann-Whitney U test. All data shown are median±interquartile range. *P*=0.0067, WT vs *Pank4*^−/−^ in 60kD group, *P*=0.1 vs *Pank4*^−/−^ in 120kD group, *P*=0.8498, WT vs *Pank4*^−/−^ in 240kD group, n=3. (**G**) and (**H**) Levels of proteins and relative quantifications of p-PKM2(Y105), p-PKM2(S37) and t-PKM2 in WT and *Pank4*^−/−^ mice lens epithelium. Data were analyzed by student’s *t* test. *P*=0.6683, WT vs *Pank4*^−/−^, n=6, in Y105 group; *P*=0.0022, WT vs *Pank4*^−/−^, n=6, in S37 group. (**I**) Immunofluorescence staining of p-PKM2(Y105) in lens epithelium of WT and *Pank4*^−/−^ mice. (**J**) Plot profile analysis described the level of p-PKM2(Y105) in WT and *Pank4*^−/−^ mice lens epithelium. (**K**) Plot profile analysis of represented single lens epithelial cell immunofluorescence stained with p-PKM2(Y105). (**L**) Immunofluorescence staining of p-PKM2(S37) in lens epithelium of WT and *Pank4*^−/−^ mice. (**M**) Plot profile analysis described the level of p-PKM2(S37) in WT and *Pank4*^−/−^ mice lens epithelium. (**N**) Plot profile analysis of represented single lens epithelial cell immunofluorescence stained with p-PKM2(S37). (**O**) Schematic illustration of current results in figure 4.

To confirm the downstream role of PKM2 that was negatively regulated by PANK4, PKM-IN-1 was used to inhibit PKM2 elevation when PANK4 was ablated. As expected, PKM2 expression was decreased but that of phosphor-PKM2 was elevated, as well as those of EMT markers, such as α-SMA, collagen I, and N-cadherin (Fig. 4C). Because PKM2 activation attenuates the lens EMT phenotype, PKM-IN-1 was injected into the anterior chamber in the TGF-β2-induced lens epithelium fibrosis model. By slit-lamp observation, an opaque anterior lens (fibrosis) was observed in the PKM-IN-1-treated group at 3 d, but a transparent lens was observed in the Veh group (Fig. 4D), which revealed PANK4 deficiency-dependent activation of PKM2.

Since we found PANK4 deficiency-related PKM2 elevation, we ask further mechanisms. A previous study indicated that cytoplasmic PKM2 functions with tetramer formation, whereas nuclear PKM2 functions with dimer/monomer formation that may provide protein-binding ability[54]. Here, we found increased expression of the 60-kDa band and 120-kDa band, corresponding to monomeric and dimeric PKM2 (Fig. 4E and F; *P*=0.0286, WT *vs. Pank4^−/−^*, respectively, n=4). Further, p-PKM2 (Y105) showed no significant difference between WT and *Pank4*^−/−^ groups, but p-PKM2 (S37) indeed declined (Fig. 4G and H; *P* =0.6683, WT *vs. Pank4^−^/^−^*, n=6, in the Y105 group; *P* =0.0022, WT *vs. Pank4^−^/^−^*, n=6, in the S37 group), which suggested that all relative increase in t-PKM2 was due to phosphorylation of the Y105 residue. To investigate the cellular location of p-PKM2 (Y105 and S37), immunofluorescence staining was performed in WT and *Pank4*^−/−^ LECs. p-PKM2 (Y105) signals were mainly located in the LEC nucleus in the WT group but in the LEC cytoplasm in the *Pank4*^−/−^ group (Fig. 4I). Plot profile of immunofluorescence signal described the p-PKM2 (Y105) level in the lens epithelium of WT and *Pank4*^−/−^ groups (Fig. 4J). To investigate the cellular location of p-PKM2 (Y105), a transection of the single cellular profile was delineated (Fig. 4K), supporting the immunofluorescence results that p-PKM2 (Y105) was mainly located in the cytoplasm and nucleus. Meanwhile, p-PKM2 (S37) immunofluorescence signals were punctate and located in the cytoplasm and nucleus of WT LECs; however, few punctate signals were found in *Pank4*^−/−^ mice (Fig. 4L). Plot profile analysis indicated lower levels of p-PKM2 (S37) optical density values in both cytoplasm and nucleus (Fig. 4M). For a single LEC, the curve indicated a typical lower level of p-PKM2 (S37) in the LEC nucleus in *Pank4*^−/−^ mice (Fig. 4N). These results indicate that PANK4 deficiency contributes to the increase in monomeric and dimeric forms of p-PKM2 (Y105) but not of p-PKM2 (S37), as indicated in the schematic diagram (Fig. 4O).

### PANK4 deficiency is responsible for PKM2 phosphorylation at Y105 and HIF-1**α** activation

PKM2 was previously reported as a PHD3-stimulated coactivator for hypoxia inducible factor-1 (HIF-1) to favor the genes associated with glycolysis[55]. Tyrosine phosphorylation of PKM2 at Y105 inhibits the formation of a PKM2 tetramer by dislodging fructose 1,6-bisphosphate (FBP) binding, which was demonstrated to be important for the Warburg effect in hypoxia and cancer[56]. Thus, we speculated the relationship between PKM2 and HIF-1 signaling may contribute to lens EMT attenuation. We performed Co-IP assay to investigate why PANK4 deletion triggered LEC to a relatively hypoxic condition. Notably, binding activity in the lens epithelium was higher in WT (Fig. 5A), which demonstrated that PKM2 (Y105) physically interacted with HIF-1α. However, no binding activity of PANK4 with HIF-1 was found in *Pank4*^−/−^ mice (Fig. 5A). To verify the glycolysis status of PANK4-deficient lens epithelium, levels of HIF-1α, HIF-1β, and HIF-2α, which are typical cellular oxygen sensors, were assessed. As expected, all three hypoxic sensors were dramatically elevated in *Pank4*^−/−^ mice (Fig. 5B; Hif-1α, *P*=0.0004, WT *vs. Pank4^−^/^−^*, n=6; Hif-1β, *P*=0.0022, WT *vs. Pank4^−^/^−^*, n=6; Hif-2α, *P*=0.0075, WT *vs. Pank4^−^/*^−^, n=6), supporting the conclusion that PANK4 was a negative regulator in the shift of OXPHOS and glycolysis. Interestingly, the lactate concentration was also increased in the lens epithelium but not in blood (Fig. 5C; *P*=0.0001, WT *vs. Pank4^−^/^−^* in the lens group, n=4). Although PANK4 is a putative kinase in the PANK family, all amniote PANK4s, including human PANK4, encode Glu138Val and Arg207Trp substitutions that might deactivate kinase activity[57]; thus, its exact role is still under consideration. To investigate the causative role and mechanisms of PANK4 in EMT, we screened the mRNAs of the lens epithelium by RNA-seq in WT and *Pank4*^−/−^ mice (n=4) and performed gene set enrichment analysis (GSEA). We found that “electron transport chain”, “respiratory electron transport”, and “oxidative phosphorylation” were enriched in the WT group, which demonstrates the potential role of PANK4 in negatively regulating glycolysis. In addition, GSEA showed an enrichment of the “Glycolysis” gene cluster in the *Pank4*^−/−^ group (Fig. 5D). To assess the transformation of OXPHOS to glycolysis by PANK4, Seahorse real-time detection was used to measure OCR and ECAR when Pank4 was knocked down by siRNA*^Pank4^*. No significant difference in basal respiration was found between the two groups; however, spare respiratory capacity and maximum respiration were decreased in the siRNA*^Pank4^* group. Non-mitochondrial oxygen consumption was dramatically increased (Fig. 5E and F; spare respiratory capacity, *P*=0.0286, siCont *vs.* siPank4, n=4; non-mitochondrial oxygen consumption, *P*=0.0286, siCont *vs.* siPank4, n=4; maximum respiration, *P*=0.0286, siCont *vs.* siPank4, n=4). The ratios of glycolytic rate, glycolytic capacity, and glycolysis reserve were all significantly increased when *Pank4* was knocked down (Fig. 5G and H; glycolytic rate, *P*=0.0317, siCont *vs.* siPank4, n=5; glycolysis capacity, *P*=0.0317, siCont *vs.* siPank4, n=5; glycolysis reserve, *P*=0.0079, siCont *vs.* siPank4, n=5). These evidences suggest the essential role of PANK4 in negatively regulating glycolysis. To verify this potential, the pyruvic acid and ATP levels were assessed in the control and siRNA*^Pank4^* groups. Dramatic increase in pyruvic acid was found when *Pank4* was knocked down, but reduction in ATP was observed compared with the siControl group (Fig. 5I and J; pyruvic acid concentration, *P*=0.0286, siCont *vs.* siPank4, n=4; concentration of ATP, *P*=0.0211, siCont *vs.* siPank4, n=4). To confirm the possible role of PANK4 in catalyzing CoA, we assessed CoA production under WT and Pank4 knockout conditions. Similarly, as previously reported[57], no significant difference in CoA was observed between the two groups in lens (Fig. S2A), suggesting a putative alternative role of PANK4 in regulating EMT. To investigate how deficiency of PANK4 contributes to the incline of glycolysis or OXPHOS, we assessed acetyl CoA carboxylase (ACC) and found an unexpected elevation (Fig. S2B). Because ACC is the key enzyme in catalyzing the step of acetyl-CoA+ATP to malonyl-CoA+ADP/Pi, and the evidence showed that the ATP level decreased in the *Pank4*^−/−^ group (Fig. 5J), we speculated the step exhaust amount of ATP.

**Figure 5.**
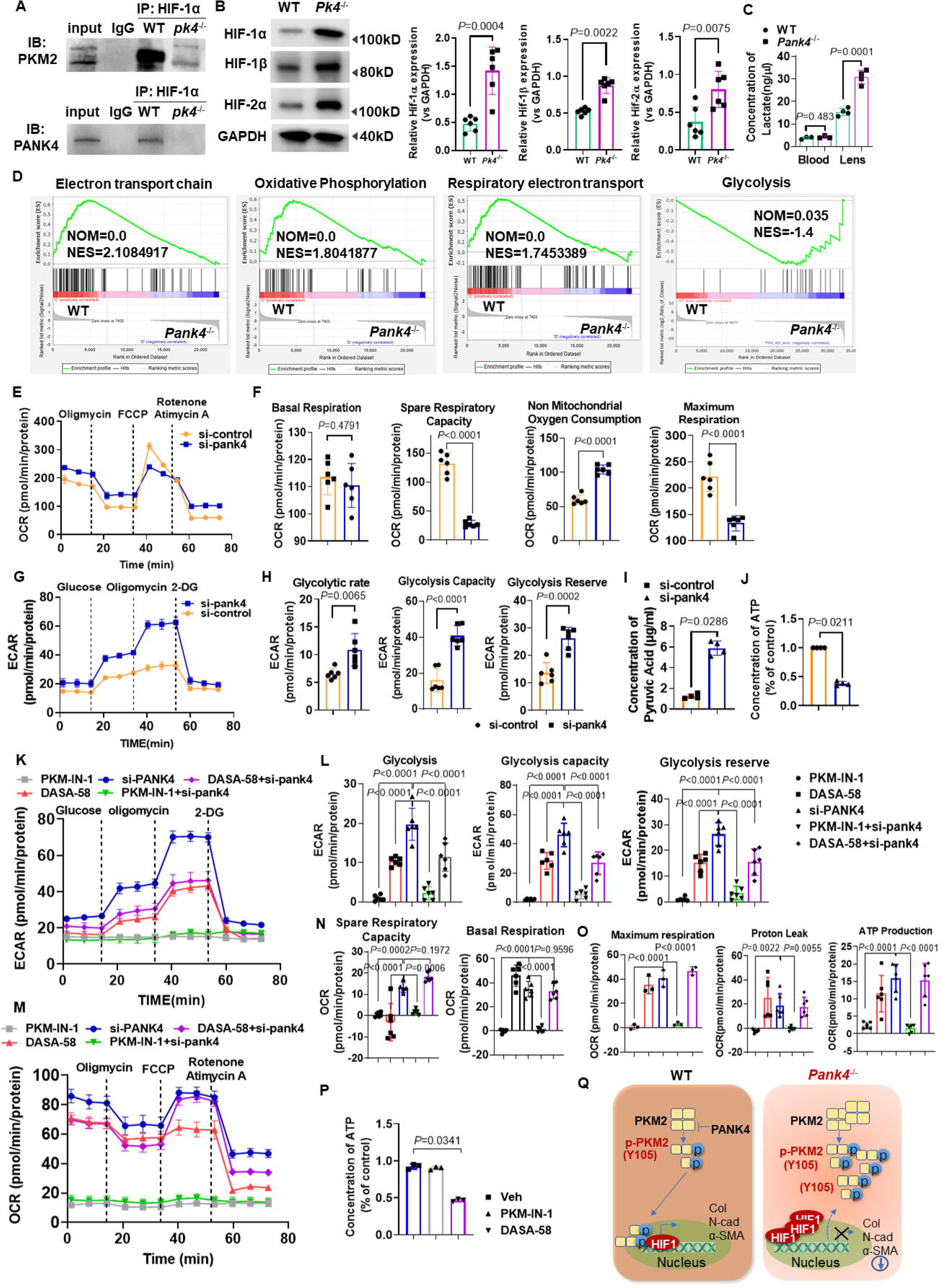
Negatively regulation of PKM2 by PANK4 in a HIF dependent way attenuating OXPHOS of lens epithelium but promoting the glycolysis related Warburg effect. (**A**) Co-IP was performed in WT and *Pank4*^−/−^ mice lens epithelium. HIF-1α was physically interacted with PKM2, but dramatically dissociated when PANK4 ablated. (**B**) The Western blot results and the quantifications indicated the protein levels of HIF-1α, HIF-1β and HIF-2α. Data were analyzed by student’s *t* test. Relative expression is the fold changes relative to the WT group. Hif-1α: *P*=0.0004, WT vs *Pank4*^−/−^, n=6; Hif-1β: *P*=0.0022, WT vs *Pank4*^−/−^, n=6; Hif-2α:*P*=0.0075, WT vs *Pank4*^−/−^, n=6. (**C**) concentrations of lactate (ng/μL) in blood and lens epithelium of WT and *Pank4*^−/−^ mice. Data were analyzed by Mann-Whitney U test. All data shown are median±interquartile range. *P*=0.0001, WT vs *Pank4*^−/−^ in lens group, n=4. (**D**) GSEA analysis enriched in electron transport chain, oxidative phosphorylation and respiratory electron transport, showed positive correlation with *Pank4*. GSEA analysis enriched in electron transport chain, oxidative phosphorylation and respiratory electron transport, showed negative correlation with *Pank4*. (**E**) and (**F**) Energy metabolism was analyzed on day 2 during reprogramming with the Seahorse instrument. Decrease in OCR after si*Pank4* was considered as the ATP production ability of the cells. Spare respiratory capacity and maximum respiration were decreased, while non mitochondrial oxygen consumption was increased after si*Pank4* treated. Data were analyzed by one way ANOVA plus Bonferroni post hoc test. Relative expression is the fold changes relative to the siControl group. Spare respiratory capacity: *P*<0.0001, siCont vs siPank4, n=6; Non-mitochondrial oxygen consumption: *P*<0.0001, siCont vs siPank4, n=6; Maxium respiration: *P*<0.0001, siCont vs siPank4, n=6. (**G**) and (**H**) Increase in extracellular acidification rate (ECAR) after si*Pank4* treatment was considered as glycolysis ability of the cells. Energy metabolism was analyzed on day 2 during Pank4 knockdown with the Seahorse instrument. Glycolytic rate, glycolysis capacity and glycolysis reserve were all increased, respectively. Data were analyzed by one way ANOVA plus Bonferroni post hoc test. Relative expression is the fold changes relative to the siControl group. Glycolytic rate, *P*=0.0065, siCont vs siPank4, n=6; Glycolysis capacity, *P*<0.0001, siCont vs siPank4, n=6; Glycolysis reserve, *P*=0.0002, siCont vs siPank4, n=6. (**I**) and (**J**) Concentrations of pyruvic acid and ATP in siControl and si*Pank4* groups in lens epithelial cells. Data were analyzed by Mann-Whitney U test. All data shown are median±interquartile range. Concentration of pyruvic acid: *P*=0.0286, siCont vs siPank4, n=4; Concentration of ATP: *P*<0.0211, siCont vs siPank4, n=4. (**K**) and (**L**) Glycolysis, glycolysis capacity and glycolysis reserve were all increased, respectively, in siPank4 group rather than PKM-IN-1, DASA-58 and PKM-IN-1+siPank4. Data were analyzed by one way ANOVA plus Bonferroni post hoc test. Relative expression is the fold changes relative to the siPank4 group. Glycolysis: *P*<0.0001, PKM-IN-1 vs siPank4, n=6; *P*<0.0001, DASA-58 vs siPank4, n=6; *P*<0.0001, PKM-IN-1+siPank4 vs siPank4, n=6; *P*<0.0001, DASA58+siPank4 vs siPank4, n=6. Glycolysis capacity: *P*<0.0001, PKM-IN-1 vs siPank4, n=6; *P*<0.0001, DASA-58 vs siPank4, n=6; *P*<0.0001, PKM-IN-1+siPank4 vs siPank4, n=6; *P*<0.0001, DASA58+siPank4 vs siPank4, n=6; Glycolysis reserve: *P*<0.0001, PKM-IN-1 vs siPank4, n=6; *P*<0.0001, DASA-58 vs siPank4, n=6; *P*<0.0001, PKM-IN-1+siPank4 vs siPank4, n=6; *P*<0.0001, DASA58+siPank4 vs siPank4, n=6. (**M**), (**N**) and (**O**) Decrease in OCR after si*Pank4* was considered as the ATP production ability of the cells. Spare respiratory capacity elevated in si*Pank4* compared with PKM-IN-1, DASA-58 and PKM-IN-1+si*Pank4* groups. Maximum respiration increased after si*Pank4* treatment compared with PKM-IN-1 and PKM-IN-1+si*Pank4* groups, while Proton leak increased after si*Pank4* treatment compared with PKM-IN-1 and PKM-IN-1+si*Pank4* groups. ATP production was elevated in si*Pank4* treatment compared with PKM-IN-1 group. Spare respiratory capacity: *P*=0.0002, PKM-IN-1 vs siPank4, n=6; *P*<0.0001, DASA58 vs siPank4, n=6; *P*=0.0006, PKM-IN-1+siPank4 vs siPank4, n=6; *P*=0.1972, siPank4 vs DASA-58+siPank4, n=6. Basal respiration: *P*<0.0001, PKM-IN-1 vs siPank4, n=6; *P*<0.0001, PKM-IN-1+siPank4 vs siPank4, n=6; *P*=0.9596, siPank4 vs DASA-58+siPank4, n=6. Maximum respiration: *P*<0.0001, PKM-IN-1 vs siPank4, n=6; *P*<0.0001, PKM-IN-1+siPank4 vs siPank4, n=6. Proton leak, *P*=0.0022, PKM-IN-1 vs siPank4, n=6; *P*=0.0055, PKM-IN-1+siPank4 vs siPank4, n=6. ATP production: *P*<0.0001, PKM-IN-1 vs siPank4, n=6; *P*<0.0001, PKM-IN-1+siPank4 vs siPank4, n=6. (**P**) Concentration of ATP (%) elevated both in Vehicle and PKM-IN-1 groups rather than that in DASA-58 group. Data were analyzed by one Kruskal-Wallis test. Relative expression is the fold changes relative to the Veh group. *P*=0.0341, Veh vs DASA-58, n=3. (**Q**) Scheme summarizing of the results.

To understand OXPHOS to glycolysis with EMT by PKM2 facilitation, Seahorse real-time detection was used to measure mitochondrial stress during the procedure (from 0 to 80 min). An increase in ECAR after adding glucose indicated the cell glycolysis level, whereas a decrease in OCR after adding oligomycin indicated the level of ATP production or OXPHOS of cells. By PKM-IN-1 treatment, glycolysis rate, glycolysis capacity and glycolysis reserve ECAR values were all decreased compared with the *Pank4* knockdown group (Fig. 5K and L; glycolysis, *P*<0.0001, PKM-IN-1+siPank4 *vs.* siPank4, n=6; glycolysis capacity, *P*<0.0001, PKM-IN-1+siPank4 *vs.* siPank4, n=6; *P*<0.0001, PKM-IN-1+siPank4 *vs.* siPank4, n=6). DASA-58 inhibited the Pank4 knockdown-related ECAR value elevation in glycolysis rate and glycolysis capacity (Fig. 5K and L; glycolysis, *P*<0.0001, DASA-58+siPank4 *vs.* siPank4, n=6; glycolysis capacity, *P*<0.0001, DASA-58+siPank4 *vs.* siPank4, n=6; *P*=0.1972, DASA-58+siPank4 *vs.* siPank4, n=6). Conversely, no difference of OCR in spare respiratory capacity, basal respiration, maximum respiration, proton leak and ATP production were found between the siPank4 and DASA-58+sipank4 groups, suggesting the glycolysis regulation of PANK4 by PKM2 (Fig. 5M, N and O; spare respiratory capacity, *P* = 0.0002, PKM-IN-1 *vs.* siPank4, n = 6; *P* = 0.1972, siPank4 *vs.* DASA-58+siPank4, n = 6; *P* = 0.0006, PKM-IN-1+siPank4 *vs.* siPank4, n = 6. Basal respiration, *P* < 0.0001, PKM-IN-1 *vs.* siPank4, n = 6; *P* = 0.9596, siPank4 *vs.* DASA-58+siPank4, n = 6. Maximum respiration, *P* < 0.0001, PKM-IN-1 *vs.* siPank4, n = 6; *P* < 0.0001, PKM-IN-1+siPank4 *vs.* siPank4, n = 6. Proton leak, *P* = 0.0022, PKM-IN-1 *vs.* siPank4, n = 6; *P*=0.0055, PKM-IN-1+siPank4 *vs.* siPank4, n =6. ATP production, *P* < 0.0001, PKM-IN-1 *vs.* siPank4, n=6). The ATP level strikingly decreased in the DASA-58 treatment group but not in the PKM-IN-1 group (Fig. 5P; *P*=0.0341, Veh *vs.* DASA-58, n = 3), which suggested that DASA-58 was effective in shifting PKM2-dependent OXPHOS and glycolysis. These results directly reflect the lower level of glycolytic activity of SRA01/04 cells by DASA-58 treatment, which could be rescued by *Pank4* knockdown. However, PKM-IN-1 did not effectively function in both ECAR and OCR analyses.

Overall, the results provide evidence that PANK4 is a regulator of EMT in lens fibrosis. PANK4 deficiency contributed to the increase in LEC glycolysis, with ATP exhaustion and thus HIFs elevation. These results indicate that PANK4 deletion may contribute to the Warburg effect by alleviating pyruvate generation. We also found a relationship between PANK4 and PKM2, in which PANK4 negatively regulated PKM2 elevation and monomer phosphorylation at Y105. Both inhibition of PANK4 and activation of PKM2 by DASA-58 resulted in EMT attenuation, which provides a potential target for lens fibrosis and PCO treatment, as indicated in the schematic diagram (Fig. 5Q).

## Discussion

Cataract is the leading cause of blindness worldwide, although it can be effectively cured by surgery. EMT is central to fibrotic cataract[58]. In the present study, PANK4, which is a PANK family member, played a vital role in LEC EMT by regulating the balance between OXPHOS and glycolysis. We also found a potential relationship between PANK4 and PKM2, and glycolysis-related HIF elevation. Consequently, we propose a model for the regulation of PKM2 in response to PANK4. PANK4 was strong enough to control LEC EMT; when it was knockout or knockdown, the expression levels of typical EMT markers, such as N-cadherin, collagen I, and α-SMA, were decreased.

PANK4 is located on chromosome 1p36.32[59] and was first described as a putative causative gene in type 2 diabetes in the Chinese population[23]. High glucose could upregulate PANK4 in rat muscle [30]. However, till now limited studies have investigated the mechanism further. Blood of diabetic patients was collected to examine the putative relationship between PANK4 and diabetes. Unfortunately, no significant difference between normal and diabetic individuals was observed in the PANK4 mRNA level (Fig. S3), which suggested that PANK4 is not a candidate causative gene for diabetes. Notably, Liu et al. reported that high PANK4 expression high PANK4 expression had shorter event-free survival and overall survival than patients with low PANK4 expression, and was indicative of poor prognosis[60], both of which need further investigation. Because of the anti-apoptotic effect of PANK4, the high level of PANK4 might inhibit pro-caspase-9 in pancreas, thus resulting in low patient survival. A *Pank4*^−/−^ mouse line was generated using the CRISPR/Cas9 system that showed an increased number of apoptotic LECs than the WT group[22]. Therefore, PANK4 may have a prominent anti-apoptotic role not only in pancreatic and myeloid cells but also in LECs.

The human genome encodes three identified PANKs (i.e., PANK1–3) plus a putative bifunctional protein, PANK4, with a predicted amino-terminal type II PANK domain and a carboxy-terminal phosphatase domain (DUF89) that is large, diverse, and present in all domains of life[20, 61, 62]. Members of the DUF89 protein family (Pfam01937) occur as C-terminal fusions with PANK in plants, animals, and chytrid fungi. Human PANK4 has activity against 4′-phosphopantetheine, and its S-sulfonate or sulfonate is comparable to that against nitrophenyl phosphate[61]. However, Yao et al. reported that all amniote PANK4s, including human PANK4, encode Glu138Val and Arg207Trp substitutions that may lack PANK activity[57]. Nonetheless, minimally characterized but highly conserved PANK4 is a rate-limiting suppressor of CoA synthesis by its metabolite phosphatase activity[63], demonstrating an essential role for PANK4 in the regulation of CoA-related metabolism. To inhibit the function of PANK4, *Pank4*^−/−^ mice that have a 5-bp mutation before exon 3 were first designed and generated. Notably, no significant difference in concentration of acetyl-CoA was found in the lens between WT and *pank4*^−/−^ mice, which demonstrated that PANK4 may be not involved in the CoA biosynthetic in lens. But we could not exclude the role of PANK4 in CoA biosynthetic in other tissues. Nonetheless, GSEA revealed a noteworthy tendency of OXPHOS, electron transport chain, respiratory electron transport, and ATP synthesis enriched in WT mice rather than in *Pank4*^−/−^ mice (Fig. 5D), suggesting the ability of PANK4 in redox regulation.

In a yeast two-hybrid system, PKM2 was associated with PANK4[30]. In the present study, we found a regulatory relationship between PANK4 and PKM2. On PANK4 loss of function, PKM2 significantly increased, suggesting a negative relationship between them, but PKM1 was unchanged (Fig. 3E). PKM2 controls the balance between energy production and metabolic synthesis precursors, and has both metabolic and non-metabolic functions, which are essential in the cytoplasm and nucleus, respectively. Tetrameric PKM2 functions as pyruvate kinase, whereas nuclear PKM2 forms a dimer and functions as protein kinase[54], which interacts with STAT3 to control downstream gene expression in SW480 cells. However, STAT3 and p-STAT3 were unchanged in the lens epithelium of *Pank4*^−/−^ mice in Western blotting test (Fig. S4), which suggested that the nuclear dimeric PKM2 regulated non-metabolic function independent of classic STAT3 signaling. PKM2 activity is inhibited after phosphotyrosine binding by the release of FBP from the PKM2 allosteric pocket by Y105 phosphorylation that disrupts tetrameric PKM2[56, 64]. Monomeric PKM2 increased under PANK4 deletion, which indicates that the glycolytic metabolic function was not likely because of tetrameric PKM2. Thus, PKM2 might not contribute to the glycolysis of LEC by tetramer formation under Pank4 ablation but is involved in p-PKM2-related signaling transduction.

PKM2-dependent histone H3 modifications are instrumental in epidermal growth factor (EGF)-induced expression of cyclin D1 and c-Myc, tumor cell proliferation, cell cycle progression, and brain tumorigenesis in its non-metabolic functions of histone modification[65]. In this study, monomeric PKM2 were increased, suggesting that the metabolic activity of PKM2 is attenuated during LEC EMT. Thus, it is conceivable that the molecular interaction of PKM2 is highly context dependent, with cell fate determined by the form of PKM2 in regulating gene expression. In addition, PKM2 plays an essential role in Warburg regulation. Methylation of PKM2 activates aerobic glycolysis to promote tumorigenesis[29]. In transformed tumor cells, PKM2 is phosphorylated at Y105 by oncogenic kinases and dissociated to PKM2 (Y105) dimers that promote yes-associated protein (YAP) nuclear translocation for tumor cell growth and transformation[66]. In this study, although PKM2 Y105 was dramatically elevated with PANK4 loss of function, the total PKM2 also increased simultaneously, demonstrating the consistent ratio of Y105 phosphorylation to total PKM2 protein, in other words, the elevated PKM2 protein was phosphorylated at Y105. Moreover, the ratio of PKM2 (S37) to total PKM2 decreased, but no significant difference was observed in the ratio of PKM2 (S37) to the inner control (Fig. 4G and H). In hepatocellular carcinoma cell lines, PKM2 could be phosphorylated at Ser37 by EGF receptor-activated extracellular signal-regulated kinase when stimulated with EGF, which promoted its nuclear translocation. By acting as both serine/threonine protein kinase and tyrosine protein kinase, PKM2 binds to and phosphorylates multiple transcription factors such as β-catenin and STAT3 in the nucleus to promote tumor-related gene transcription. H3-Thr11 phosphorylation is dependent on the phosphorylation of PKM2 at Ser37[67]. Herein, to clarify whether PKM2 (S37) phosphorylates STAT3 under PANK4 deletion, STAT3 and p-STAT3 were both detected, but no change was observed (Fig. S4). Immunostaining of PKM2 (S37) in the lens epithelium was attenuated in the *Pank4*^−/−^ group, which strengthens the conclusion that S37 residue-related PKM2 transnuclear function may not be involved in PANK4-unlocked PKM2 elevation.

The lens is sustained in a hypoxic environment in the eye without any vascular supply. The oxygen partial pressure at the lens surface is 2.9 mmHg, which is lower than 24.1 mmHg at the surface of the eye[68]. Thus, PKM2 activity contributes to a high glycolytic potential to support lens metabolism[69–71]. PKM2 can catalyze the final rate-limiting step of glycolysis, and it functions as a dimer when translocating to the nucleus, where it stimulates HIF-1α transactivation domain function and recruitment of p300 to hypoxia response[55]. In the present study, we found that HIF-1α and HIF-1β increased dramatically in parallel with total PKM2 and PKM2 (Y105) elevation under PANK4 deletion. The current results showed a physical relationship between PKM2 and HIF-1α in WT mice but not in *Pank4*^−/−^ mice. Because the lens epithelium was in a hypoxic environment in the WT condition, PKM2 and HIF-1α might contribute to the homeostasis of LEC survival and the balance of metabolism. After PANK4 was absent, more PKM2 was phosphorylated at Y105 and dissociated with HIF-1α, along with favoring glycolysis, indicating a negative regulation loop of PANK4 on hypoxia. Although *Pank4*^−/−^ mice were housed in a high-level oxygen (70%) condition for 7 continuous days, HIF-1α was still elevated in the lens epithelium of *Pank4*^−/−^ mice (Fig. S5, *P*=0.0021, WT(O2) vs *Pank4*^−/−^(O2), n=4), which suggested that HIF-1α elevation is caused not only by hypoxic conditions but also by PANK4 regulation. In normoxic conditions, HIF-1α can be hydroxylated in its C-terminal transcription activation domain by the arginyl hydroxylase factor-inhibiting HIF[72]. Oxygen-dependent HIF-1α degradation is mediated by the hydroxylation of prolines 402 and 564 in human HIF-1α. However, the Warburg effect can activate PI3K and Akt signaling to stabilize HIF-1α in an aerobic condition, which promotes pyruvate-related glycolysis as well[73]. Therefore, the consistent elevation of HIF-1α is dependent on PANK4 deletion, irrespective of normoxic, hypoxic, or hyperoxic conditions. Concerning the uncoupling result of PKM2 and HIF-1α with PANK4 loss of function (Fig. 5A), we speculate that the elevation of PKM2 and HIF-1α may function in different ways rather than the classic PKM2-HIF-1-positive feedback loop. PANK4 may function as a coordinator that organizes PKM2 and HIF-1α in cellular OXPHOS and glycolysis activities. In addition, we cannot exclude the possibility that HIF-1α promotes PKM2 gene transcription or PKM2 promotes HIF-1α gene transcription, which needs evidence. In older animals, HIF-1α is necessary to maintain a high level of p27KIP1 that inhibits LEC proliferation. Because we previously found an apoptotic phenotype of LECs in *Pank4*^−/−^ mice, the conclusion that PANK4 deficiency-dependent HIF-1α elevation may inhibit the proliferation and EMT of LECs seems plausible.

## Acknowledgement

We thank those heroes combating the 2019-nCov, especially Miss Ying Li and Juan Yu in our department, with respect. We also thank Prof. Lei Li from Shanghai Tech University for data evaluation. We thank our team members (Especially Jiao Hu, Xi Chen and Qiu-Mei Hu) for their diligent work. We also thank Cyagen Inc. for mice embryo cryopreservation and breeding. The project is supported by National Natural Science Foundation of China (No. 82070945), Science and Technology Innovation Enhancement Project of Army Medical Center, Army Medical University (No. 2019CXJSB009 and No. 2019CXJSB021), Army Medical University Outstanding Talent (Chun-Lin Chen) and grants from Basic Research and Scientific Frontier Foundation of Chongqing (cstc2019jcyj-msxmX0010).

## Supplementary Figure Legends

**Figure 1.** Screen assay of 3 types of designed siPank4 by Western blot.

**Figure 2.** (**A**) Concentrations of acetyl-CoA in WT and *Pank4*^−/−^ mice lens epithelium, brain cortex and retina. Data were analyzed by Mann-Whitney U test. All data shown are median±interquartile range. Lens: *P*=0.7, WT vs *Pank4*^−/−^, n=3; Retina: *P*=0.1, WT vs *Pank4*^−/−^, n=3; Brain: *P*=0.4, WT vs *Pank4*^−/−^, n=3. (**B**) The protein level and quantification of acetyl CoA carboxylase (ACC) in WT and *Pank4*^−/−^ lens. Data were analyzed by Mann-Whitney U test. All data shown are median±interquartile range. *P*=0.0286, WT vs *Pank4*^−/−^, n=4.

**Figure 3.** Relative mRNA of Pank4 in healthy and diabetes patients. Data were analyzed by Mann-Whitney U test. All data shown are median±interquartile range. *P*=0.8413, WT vs *Pank4*^−/−^, n=5.

**Figure 4.** Protein levels of total-Stat3 and phosphor-stat3 in WT and *Pank4*^−/−^ mice lens epithelium. Data were analyzed by Mann-Whitney U test. All data shown are median±interquartile range. *P*>0.9999, WT vs *Pank4*^−/−^, n=3.

**Figure 5.** Protein levels of PKM2 and HIF-1α in WT and *Pank4*^−/−^ mice lens with and without 70% oxygen treatment for continuous 7 days. Data were analyzed by Mann-Whitney U test. All data shown are median±interquartile range. *P*=0.0286, WT *vs. Pank4*^−/−^, n=4; *P*=0.0286, WT(O2) *vs. Pank4*^−/−^(O2), n=4.

