## Supplementary figures and images for "Pantothenate Kinase 4 Governs Lens Epithelial Fibrosis by Negatively Regulating Pyruvate Kinase M2-Related Glycolysis"

### suppl fig 1

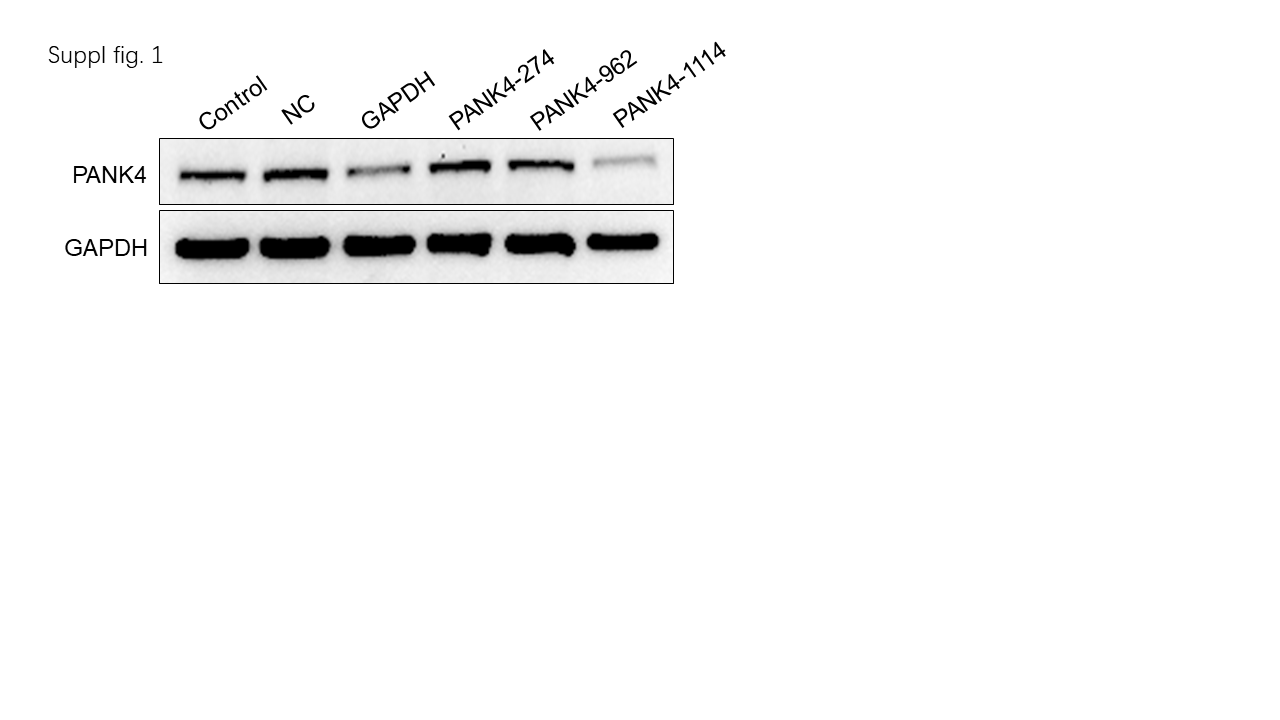

### suppl fig 2

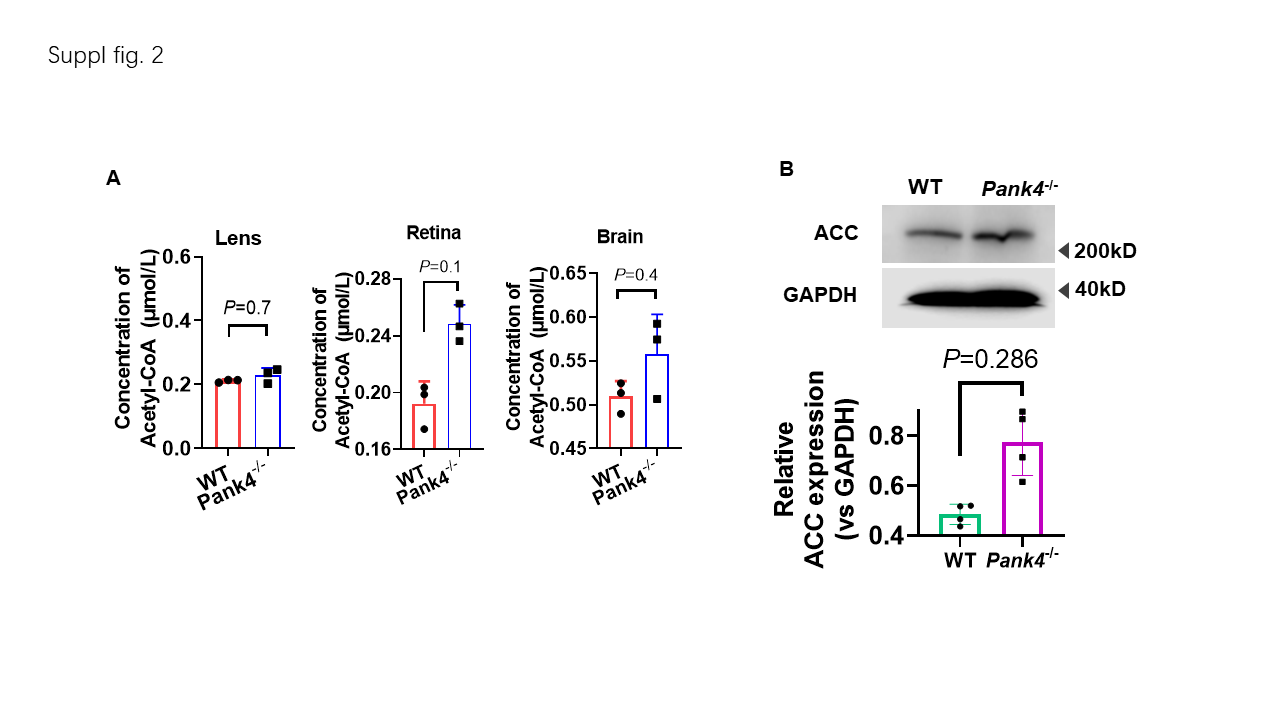

### suppl fig 3

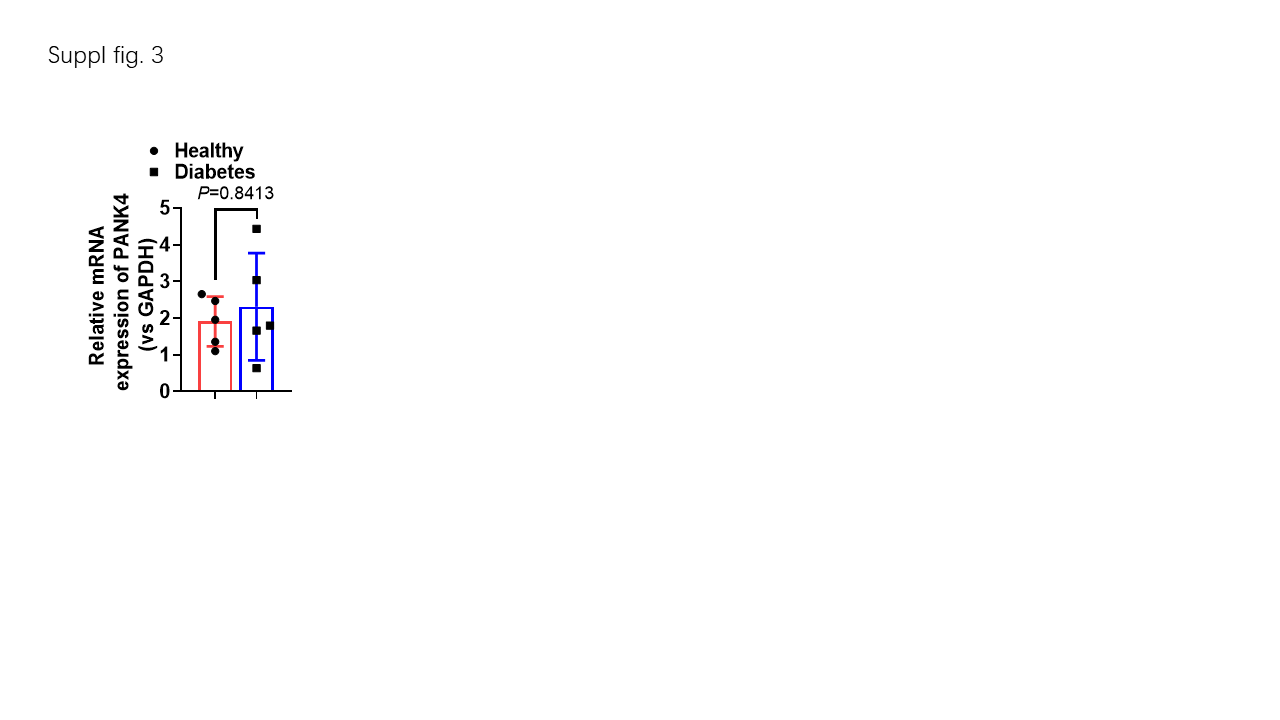

### suppl fig 4

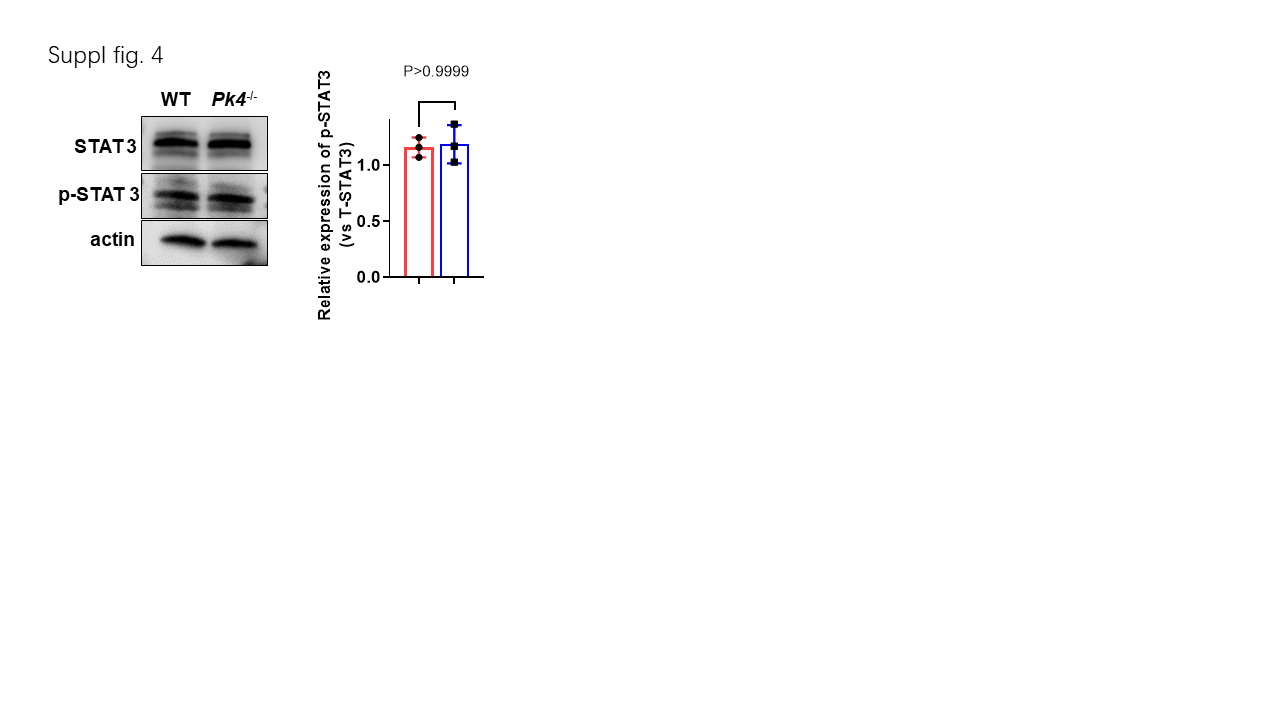

### suppl fig 5

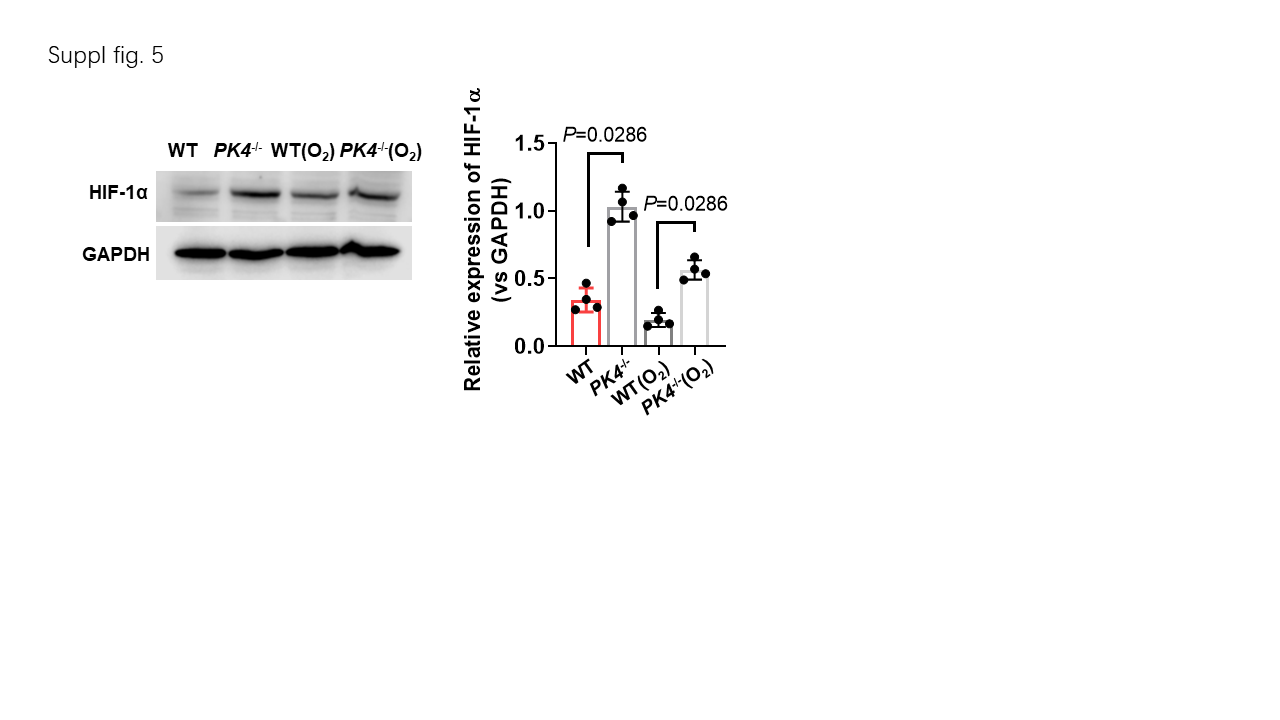
